# Behavioral Decoding Reveals Cortical Endocannabinoid Potentiation during Δ^9^-THC Impairment

**DOI:** 10.1101/2025.09.26.678874

**Authors:** Anthony English, David Marcus, Khushi Yadav, Yassin Elkhouly, Allan Levy, Victoria Corbit, Maddie Ask, Anika Scedberg, Jordan Poces-Ball, Fleur Uittenbogaard, Rayna Simons, Ilana Witten, Larry Zweifel, Benjamin Land, Nephi Stella, Michael R. Bruchas

## Abstract

How Δ^9^-tetrahydrocannabinol (THC) impairs natural behaviors in mice remains unknown. We developed a video-monitored behavioral platform with machine learning classifiers to unravel discrete changes in natural mouse behaviors. THC infusion into the medial prefrontal cortex (mPFC) disrupted walking kinematic features characteristic of impairment responses. THC predominantly increased mPFC GABAergic activity preceding walk initiation shifting the mPFC excitatory/inhibitory (E/I) balance. Pose-defined closed loop photo-stimulation of mPFC GABAergic neurons demonstrated that THC exacerbates selected parameters of motor impairment. Surprisingly, THC also induced a time locked, movement-induced, transient potentiation of mPFC endocannabinoid (eCB) release and ensuing CB_1_R-mediated synaptic inhibition. Here we establish that THC-modifies mPFC E/I balance to excitation via dynamic changes in eCB release which acts to induce behavioral impairment.

## 1 Introduction

The increasing use of Cannabis-based products that contain high amounts of THC has raised concerns about their acute effects on natural behaviors, particularly on spontaneous locomotion and its kinematics and coordination (1, 2). While THC’s ability to reduce spontaneous locomotion in mice monitored in an open-field setting is well-recognized, our fundamental understanding of THC’s bioactivity on influencing naturalistic behaviors has been limited by current experimental approaches (3–5). Specifically, genetic and molecular approaches have established that THC modulates neuronal activity by engaging presynaptic CB_1_Rs (6). Numerous rodent studies have demonstrated that high doses of THC reduces the overall speed of locomotion in mice through CB_1_R activation localized to the cortex and forebrain (7, 8). The underlying impairing effect of THC on specific features of naturalistic locomotor behavior, including movement initiation and walking gait parameters are unknown, critical knowledge important for understanding THC induced impairments in humans.

The mPFC plays a fundamental role in movement planning, including the preparation for spontaneous locomotion and adaptive action timing (9–11). The mPFC expresses one of the highest densities of CB_1_R in brain, implicating its involvement in THC’s impact on natural movement releated, motoric behaviors (12). Prior studies investigating THC’s effects on inhibiting spontaneous locomotion and inducing catalepsy used monitoring approaches that lack the spatiotemporal resolution required to uncover disrupted naturalistic behavioral signatures and whether they are specifically coupled to changes in neural activity during THC impairment (11, 13–16). In parallel, the development of biosensors that detect changes in neural activity and subcellular changes in eCB levels now provide unprecedented means to measure real time dynamics of psychoactive drugs on neural circuits during discrete behavioral (17, 18).

Computer vision tools, such as DeepLabCut and SLEAP, have begun to transform current behavioral research by allowing for high-resolution tracking of individual points on animals during expression of behavioral phenotypes (19–22). These tools have substantially improved our ability to define uniquely segmented behavioral events; yet they have not been utilized within the neuropharmacology landscape coupled to neural circuit recordings, including the defining of drug-induced behavioral action. Here we designed a SLEAP pose estimation algorithm and analysis pipeline for spontaneous and naturalistic behavior in freely moving mice within a two-camera view linear track system (side and bottom). Using this approach, we isolated the impact of acute THC treatment on distinct naturalistic mouse behaviors identified frame by frame. We then applied a computational approach wherein THC-specific dose prediction algorithms decode the “presented behavioral dose” of control and genetically manipulated mice. We concomitantly measured changes in mPFC neural activity and eCB levels using cell type-specific expression of the biosensors to monitor THC-induced shifts in E/I balance within the mPFC. Predictive behavioral impairment algorithms were then applied in concert with online closedloop optogenetic approaches to actuate real-time mPFC neural activity during THC-induced impaired movement. The results presented here uncover a mechanism whereby THC-induced impairment of spontaneous locomotion shifts E/I balance and reveal specific epochs of potentiated eCB release within the mPFC through actions at CB_1_R. By integrating behavioral, neural, and computational measures, our approach provides a lens into the mechanisms of THC-induced impairment, to predict treatment dose and uncover the signatures driving disrupted behavior during THC intoxication.

## 2 Results

### Detection of THC-induced impairment signatures using machine learning and dose-prediction computational models

THC-induced locomotor impairment has been traditionally quantified in rodents by calculating total distance traveled or average velocity over time in an open field arena as measured by beam breaks (23, 24). Such THC-induced reduction in overall spontaneous locomotion is included in the canonical “tetrad” response (hypolocomotion, catatonia, hypothermia, and analgesia) developed as an initial industry standard to measure CB_1_R-dependent cannabimimetic bioactivity (25, 26). This general decrease in spontaneous locomotion fails to identify changes in discrete natural behaviors induced by THC while mice are moving. To address this limitation, we implemented and trained a pose estimation algorithm to monitor mice spontaneously moving in a custom built linear track corridor that leverages a mirror positioned below at a 45*°* angle for dual-viewpoints (i.e., side and bottom-up) using a single, high frame-rate camera (Figure 1A) (27, 28). To detect distinct natural, spontaneous behavioral events with higher resolution, we developed an AI pose estimation algorithm that monitors 30 precise points detected from both views. The algorithm was trained with over 1,700 hand-labeled frames from 72 videos of the naturalistic behavior of WT male and female mice freely moving in the chamber Supplemental Figure S1). A secondary random forest (**RF**) supervised classification algorithm was developed to automatically classify 4 core behaviors (walking, rearing, grooming, and laying on belly (**LOB**)) with *>* 95% accuracy when trained on 176 videos (1.6 million frames) of hand-labeled data (Figure 1A and Supplemental Figure S1). Mice were treated with increasing doses of THC and 1 h later (i.e., a time point known to result in strong mouse brain exposure to THC (29) placed in the linear track for unbiased recording. Figure 1B shows a behavioral ethogram exemplifying the high temporal discretion of positional classification over the course of the first 5 min analysis in the linear track. THC-induced a dose-dependent reduction in grooming (IC_50_ = 0.10 mg/kg), rearing (IC_50_ = 0.62mg/kg) and walking (IC_50_ = 0.83mg/kg) and increased LOB (EC_50_ = 1.23 mg/kg) (Figure 1C). Classified walk events were isolated, and point-wise kinematic analyses revealed distinct effects of THC on walking behavior (Figure 1D and Supplemental Figure S2). Motor kinematics, such as gait width, are a known indicator of motor instability (30). Of the specific kinematic parameters, forepaw gait width and hind paw gait width (i.e., the average widths of the paws over the walk event) were sensitive to THC as indicated by their dose-dependent increase, indicating that our approach can detect subtle instability by motor kinematic impairment induced by THC in a dose-dependent manner (Supplemental Figure S2). We then standardized an array of kinematic metrics across THC doses to vehicle control to isolate THC’s pharmacodynamic profile as it relates to specific kinematic metrics during walking (Figure 1E).

**Figure 1.**
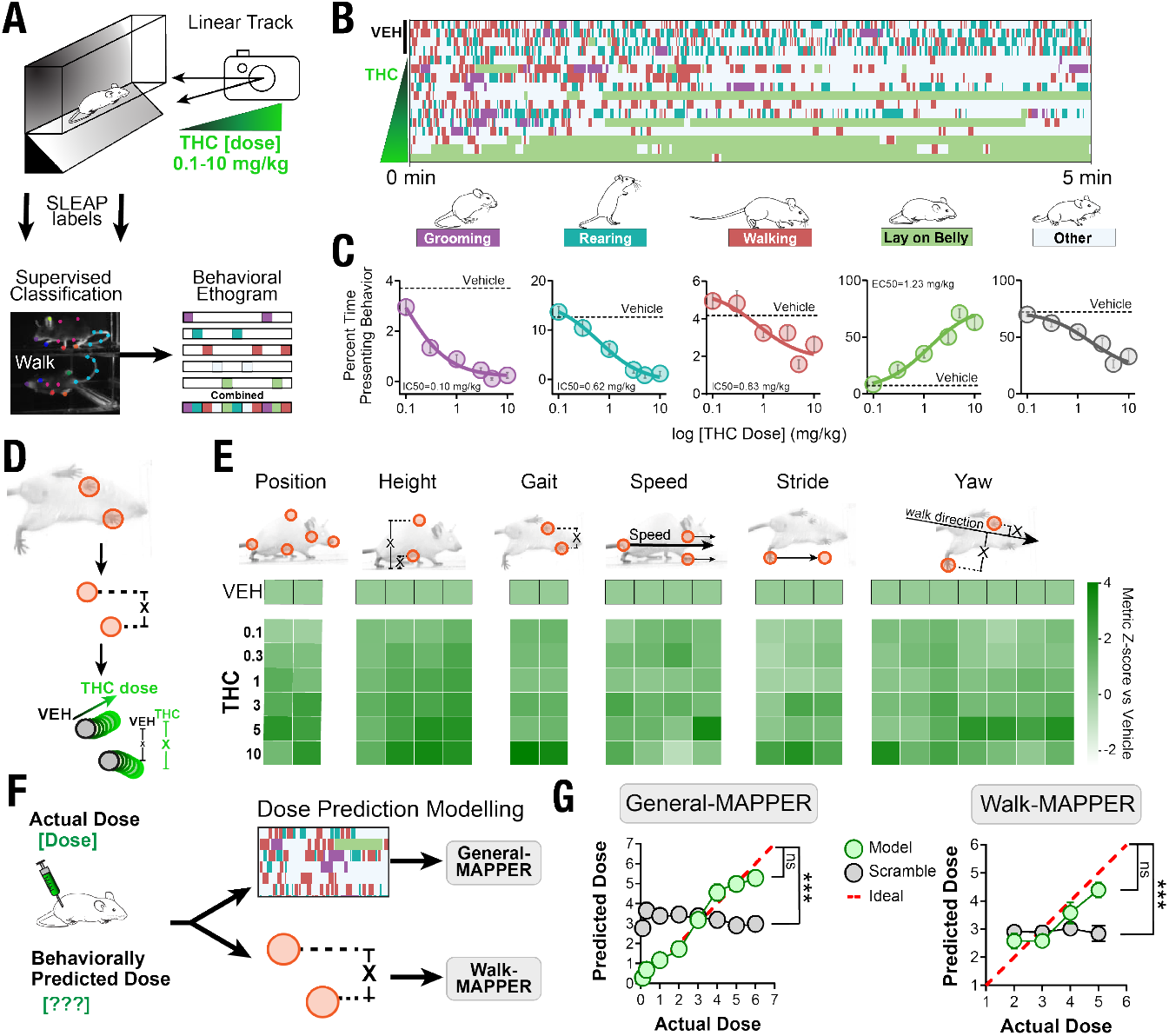
Detection of THC-induced impairment signatures using machine learning and dose-prediction computational models: **A)** Schematic of analysis pipeline for linear track recordings first analyzed with a supervised classification algorithm. **B)** Behavioral ethogram over 5 minutes in the linear track identifying core behaviors (grooming, rearing, walking, and laying on belly. Each row represented one mouse treated with vehicle or increasing doses of THC. **C)** THC dose-dependent response curves for percent time in four core behaviors as assessed by supervised classification algorithm after a two-day habituation period. **D)** Schematic of kinematic analysis as calculated by POIs during walk events. **E)** Kinematic metrics during walk events after increasing doses of THC with Zscores standardized to vehicle treatment. **F)** Schematic for training random forest regression models to predict THC dose from general behaviors and from kinematic features. **G)** Dose prediction accuracy of general and kinematic behavioral models against a test set (green) and a dataset scrambled across doses (grey) compared to an ideal prediction (red). Nonlinear fit curve significance from ideal, ^***^p<0.0001. Full experimental dataset: vehicle (N=179) and THC (0.1-10 mg/kg: N = 10-35 each). Random forest regressor training set: THC (0.1-5 mg/kg: N = 16-35 each).

Leveraging the high tracking accuracy of our approach, we constructed a non-linear RF regression model trained by n = 364 videos of the dose-dependent effect of THC to predict the “presented dose” of THC on each core natural behaviors (walking, rearing, grooming, and LOB) (Figure 1F). Furthermore, we leveraged the kinematic metrics calculated in Figure 1E as a new set of features to train a second dose prediction model designed to predict the dose of THC that impairs kinematic motor properties during walk events (Figure 1F). This model serves to identify the severity of motor impairment rather than the general THC-effects captured in the first model. As a control, we presented to these models selected videos of mice treated with a range of THC doses and shuffled the identified behavioral signatures to eliminate the algorithm’s dose prediction accuracy. Figure 1G shows that the developed dose prediction algorithms are accurate while the shuffled was significantly different from the “ideal” prediction. These results establish that our dose prediction models decode the effective dose of THC treatment based solely on computationally visualized behavioral data sets. Of note, both models displayed their nearest to ideal accuracy in the effective THC dose range between 2 and 5 mg/kg (31–33). Together, these results indicate that the two models perform the supervised monitoring of naturalistic behaviors allowing for precise quantification and dose-prediction of drug treatment.

### Unsupervised modeling defines unique changes in behavioral signatures induced by THC

Across all treatment groups, including vehicle and THC at all doses, 26.4% of naturalistic behaviors were not identified as walking, rearing, grooming, and LOB, and thus categorized as “Other,” indicating that additional naturalistic behaviors remained to be identified. To further quantify the complexity of THC’s impact on mouse behavior, we used an unsupervised, high-dimensional K-means clustering algorithm to isolate previously unrecognized behavioral patterns (Figure 2A). We excluded all frames classified as walking, rearing, grooming, or LOB (“other” frames) in 432 videos of THC-treated mice. Figure 2B shows 15 discrete behavioral clusters identified in the remaining “other” frames when analyzing a set of 29 positional features using a 29-dimensional K-means clustering framework. Specifically, the centroid point of each of the 15 identified clusters and its size is scaled to reflect the relative quantity of frames represented within each cluster (Figure 2B). Thus, Figure 2B shows that the plots size and distributions depicting these other mouse naturalistic behaviors were significantly shifted by 5 mg/kg THC treatment compared to vehicle. Figure 2C shows the THC dose-response curves for each cluster revealing that this treatment induced specific effects characterized by a decreased and increase in their occurrence, or that were unaffected by THC as compared to controls (Supplemental Figure S3). For example, THC-induced a robust decrease in the frequency of a crouching position (post-hoc here named “crouching”) (Figure 2D). The ability of unsupervised clustering analysis to identify THC-induced changes in the occurrence of these behavioral signatures emphasizes the value of this “black box” un-biased classification approach based on neuropharmacological datasets to identify behavioral readouts not detectable by human observations. Similar to the dose prediction models in Figure 1, calculated features were utilized in an unsupervised dose prediction model. These features were combined with those from the core prediction model to then train a “comprehensive dose prediction” algorithm, which reached a mean square error (**MSE**) of 0.47, surpassing the core model alone (compare Figure 2E and Figure 1E). The high accuracy of the Comprehensive Prediction Model demonstrates the incorporation of unsupervised clustering in feature detection for unique THC-induced impairment signatures.

**Figure 2.**
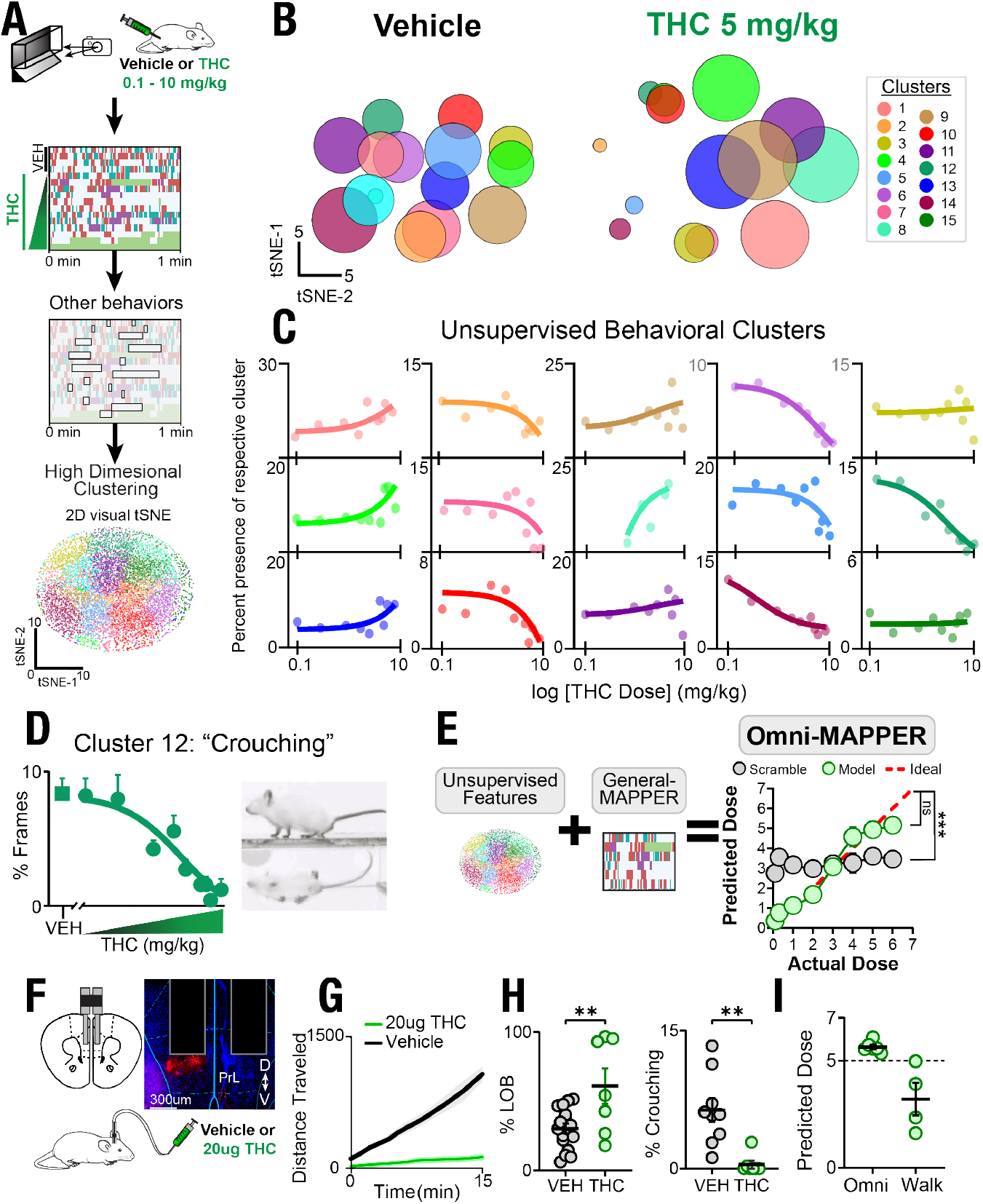
Unsupervised models reveal unique behavioral signatures of THC impairment mediated by mPFC activity: **A)** Schematic of analysis pipeline for extracting nuanced “other” frames to generate features for high dimensional Kmeans clustering (N=179, 53% male, 47% female, treatment range vehicle, 0.3, 0.1, 1, 2, 3, 4, 5, 6, 7, or 10 mg/k THC). 2 dimensional (2D) tSNE shows representative segmentation of clusters. **B)** To visualize structure within the high-dimensional dataset, we applied a two-dimensional t-SNE to the 29-dimensional behavioral feature space. Vehicle and 5 mg/kg THC treated behavioral frames with unsupervised cluster centroids sized to relative contribution for given treatment. **C)** Relative expression per “other” contributions were plotted and THC dosedependent effects of 16 identified clusters. **D)** Crouching “crouching” dose response curve along with a visual snapshot of behavior and 2D tSNE with cluster highlighted. **E)** Schematic for training of a random forest regressor for dose prediction by combining an unsupervised model using only nuanced behavioral clusters and the combination with the general model to train a Comprehensive behavioral dose prediction model. Nonlinear fit curve significance from ideal, ^***^p<0.0001. **F)** bi-lateral intracranial infusion of high concentration THC (10 ug each side). **G)** Standard behavioral effect of total distance traveled in linear track. **H)** Percent laying on belly and expression of Crouching exemplifies THC effect. **I)** Application of Comprehensive and kinematic dose prediction models reveals high dose THC behavioral presentation. N=7-15 Student’s unpaired T-test (^**^p<0.01).

### THC acts within the mPFC to induce signatures of behavioral impairment

Considering the role of the mPFC in the control of movements, we implanted cannulas into the prelimbic cortex of mice (bilateral: AP = 1.8, DV = − 2.0, ML = 0.4), infused bilaterally either vehicle or THC (10 µg/1 µl) and recorded their naturalistic behavior in the linear track (Figure 2F) (34, 35). THC infusion in the prelimbic cortex was sufficient to reduce locomotion measured as distance traveled (Figure 2G) (36). Furthermore, analysis with the supervised classification algorithm indicated a significant increase in time spent LOB, and a significant decrease in the crouching behavior following THC infusion (Figure 2H). Application of the Comprehensive and Kinematic dose prediction models predicted that local mPFC infusion induced standard THC-like behaviors with a predicted dose of 5.64 mg/kg and 3.21 mg/kg, respectively (Figure 2I). The underprediction by the Kinematic model to assess locomotor impairment at a 3.2 mg/kg compared to the Comprehensive model indicates an important role for the mPFC in mediating THC’s locomotor impairing effects.

### THC shifts E/I balance to excitation in the mPFC during walking epochs

To identify the neural circuit within the mPFC during THC-induced impairment, we expressed the fluorescent calcium indicator, GCaMP6f, in mPFC^CAMKII*α*^ neurons to monitor changes in the activity of these projection neurons (Figure 3A). Specifically, mPFC^CAMKII*α*^ (AAV-DJ-CAMKII*α*-GCaMP6f) mice were treated with vehicle or THC (5 mg/kg, *i*.*p*.) and 1 h later monitored for 15 min in the linear track coupled to simultaneous fiber photometry recordings. We found that the activity of projection neurons was time-locked to the initiation of each walk event in THC-treated mice (Figure 3B). Averaging neural activity across all walk events of THC-treated mice revealed a ramping increase in GCaMP fluorescence that preceded a walk event, which persisted for the length of a walk event (1.8*±*1.4 s) and then returned to baseline (Figure 3B). This response also occurred in mice treated with higher doses of THC (peak Z-score = 2.73*±*0.03) and was absent in vehicle-treated animals (peak Z-score = 0.24 *±* 0.01).

**Figure 3.**
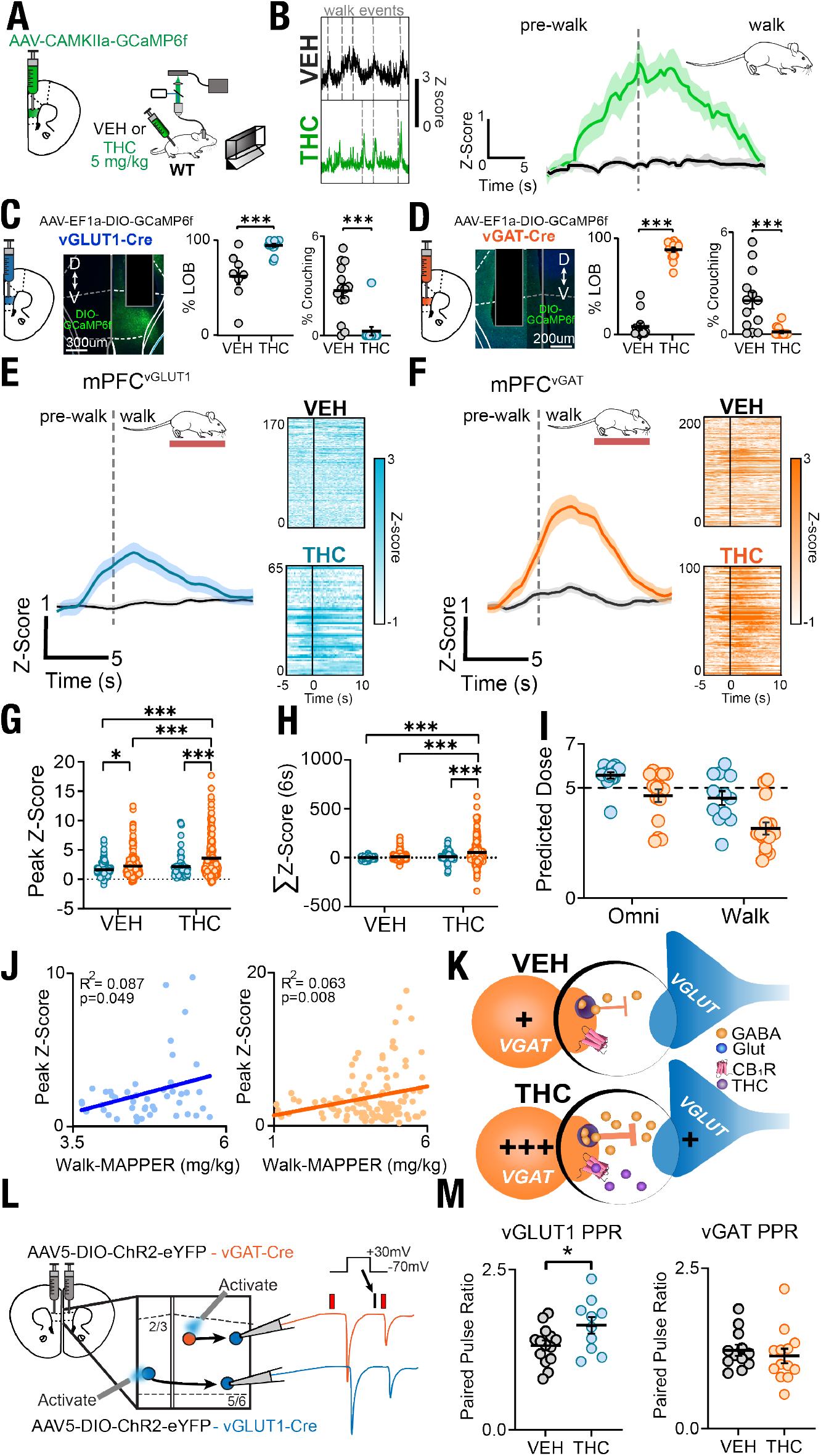
THC modifies E/I balance in the mPFC during locomotor walk epochs: **A)** mPFC injection of AAV-DJ-CAMKII*α*-GCaMP6f for fiber photometry linear track experimentation. **B)** Time locked mPFCCAMKII*α* neural signal at walk initiation during WT-CAMKII behavior. Averaged walk event traces after vehicle or 5 mg/kg THC treatment. **C-D)** Schematic and histology for Cre-dependent expression of GCaMP6f in mPFC^vGLUT1^ (**A**) and mPFCvGAT (**B**) neurons. Percent LOB and Crouching behavioral presentation in vehicle and 5 mg/kg THC-treated mPFC^vGLUT1^ neurons (**C**) and mPFC^vGAT^ (**D**) neurons. **E-F)** Fiber photometry signal of glutamatergic (**E**) and GABAergic (**F**) neuron activity at walk initiation and subsequent heatmaps for each walk event. **G-H)** Peak Z-score (**G**) and Σ Z-score (6 s) (**H**) over neural recordings of walk events in mPFC^vGLUT1-GCaMP6f^ and mPFC^vGAT-GCaMP6f^ mice. **I)** Application of Comprehensive and Kinematic dose prediction models to VGLUT-GCaMP6f behavior during fiber photometry recording. **J)** Application of Comprehensive and Kinematic dose prediction models to VGAT-GCaMP6f behavior during fiber photometry recording. **K)** Cartoon schematic of mPFC neurons after THC administration during walk behavior. **L)** Schematic of electrophysiological recordings of mPFC^vGLUT1^ and mPFC^vGAT^ neurons. M) Normalized oEPSC amplitude after optogenetic stimulation with and without THC bath reveals enhanced maximum DSE in mPFC^vGAT^ neurons. N=8-14, Two-Way ANOVA with Sidak’s posttest and Student’s unpaired T-test (^*^P<0.05, ^**^p<0.01, ^***^p<0.001).

We next used FISH to map CAMKII*α* expression to better isolate THC’s effect on E/I balance and found that CAMKII*α* was expressed in both GABAergic and glutamatergic neurons within the mPFC (Supplemental Figure S4). Because of this non-selectivity we used a cre-driver approach to directly assess the activity of glutamatergic and GABAergic neurons with vGLUT1-Cre (mPFCvGLUT1) and vGAT-Cre (mPFCvGAT) mice injected with AAV5-DJ-EF1a-DIO-GCaMP6f, respectively, and implanted with an optic fiber within the mPFC (Figure 3C-D). Mice were treated with either vehicle or THC (5 mg/kg, *i*.*p*.) and 1 h later monitored in the linear track for 15 min. As expected, THC increased the percent time spent LOB and decreased crouching in both cre-driver lines compared to vehicle treatment (Figure 3C-D). We then isolated walk events using the supervised algorithm to time-lock changes in mPFC neural activity to walk initiation. THC induced an increase in mPFC^vGLUT1^ neural activity that began to increase 3-4 sec before movement onset and reached an average peak Z-score of 1.53 *±* 0.02 (Figure 3E). Increased mPFC^vGAT^ neural activity was similarly time-locked to walk initiation, began 3-4 sec prior to walk initiation, and reached a higher peak activity (Z-score = 2.51 *±* 0.03) after THC treatment compared to vehicle (Z-score of 0.82 *±* 0.01), indicating a more pronounced increase in inhibitory activity (Figure 3F). The increased GABAergic activity, time-locked to THC-dependent behavior, was CB_1_R-dependent because pre-treatment with the competitive antagonist SR141716A (**SR1**) blocked THC-induced transient activity (Supplemental Figure S5). Further analysis indicated that THC-treated mPFC^vGLUT1^ and mPFC^vGAT^ neurons showed significantly higher cumulative Z-score for the 1 second before and the 5 seconds after walk initiation (Σ Z-scores (6 s)) compared to vehicle (Figure 3G-H). Application of the Comprehensive Prediction Model (“Com”) revealed a presented behavioral dose of 5.58 *±* 0.14 mg/kg and 4.62 *±* 0.29 mg/kg collected from mPFC^vGLUT1-GCaMP6f^ and mPFC^vGAT-GCaMP6f^ mice showing that the experimental setup did not obscure THC’s gross cannabimimetic effects (Figure 3I). Kinematic Dose Prediction (“Kine”) revealed a difference between THC-treated mPFC^vGLUT1^ and mPFC^vGAT^ animals, suggesting a potential underlying strain-dependent effect. We next isolated peak Z-score activity detected in mPFC^vGLUT1^ and mPFC^vGAT^ neurons and the kinematic dose prediction for each individual walk event. This analysis revealed a significant correlative relationship in both populations (mPFC^vGLUT1^: p=0.049 and mPFCvGAT: p=0.008) (Figure 3J). These results indicate that THC modifies the activity of both mPFC^vGLUT1^ and mPFC^vGAT^ neurons, ultimately shifting the E/I balance during walking epochs, to potentially promote impairment.

To better understand this change, we tested if the E/I shifts could be acting in a compensatory, homeostatic manner to correct the impaired movement wherein enhanced GABA release modulates mPFC^vGLUT1^ tone (Figure 3K). Therefore, we used slice electrophysiology to record paired pulse ratio (PPR) in mPFC^vGLUT1^ and mPFC^vGAT^. Specifically, mPFC^vGLUT1^ and mPFC^vGAT^ neurons were selected via injection with the light-activated opsin ChannelRhodopsin (AAV5-DIO-ChR2-eYFP) into either the contralateral or ipsilateral mPFC, respectively, to pinpoint the ex vivo slice electrophysiology recordings (Figure 3L and Supplemental Figure S6). Tissues were bath-treated for 1 h in either vehicle or THC (20 *µ*M). THC-induced an increase in Paired Pulse Ratio (**PPR**) from mPFC^vGLUT1^ neurons as indicated by a reduced excitatory input to pyramidal neurons and an unchanged inhibitory input, confirming a THC-induced change in the E/I balance to excitation (Figure 3L-M). Together, these results provide evidence for a THC-dependent shift from excitation to inhibition within the mPFC that influencing E/I balance and naturalistic walking behavior.

### CB_1_R Signaling within mPFC GABAergic Neurons Underlies THC-Driven E/I Imbalance

To determine whether CB_1_R mediates the modulatory effects of THC within the mPFC, and to isolate the specific neuronal population involved in the THC/CB_1_R-induced impairment of specific kinematic parameters during locomotor events, we utilized a conditional CRISPR-saCas9 CB_1_R knockout in either mPFC^vGLUT1^ or mPFC^vGAT^ neurons. Specifically, we bi-laterally injected either CB_1_R CRISPR (AAV1-FLEX-saCas9-sgCnr1) or ROSA control (AAV1-FLEX-saCas9-sgROSA26) viruses into the mPFC (600 nl: AP: 1.8, DV: -2.0, ML: *±*0.4) (Figure 4A) (37).

**Figure 4.**
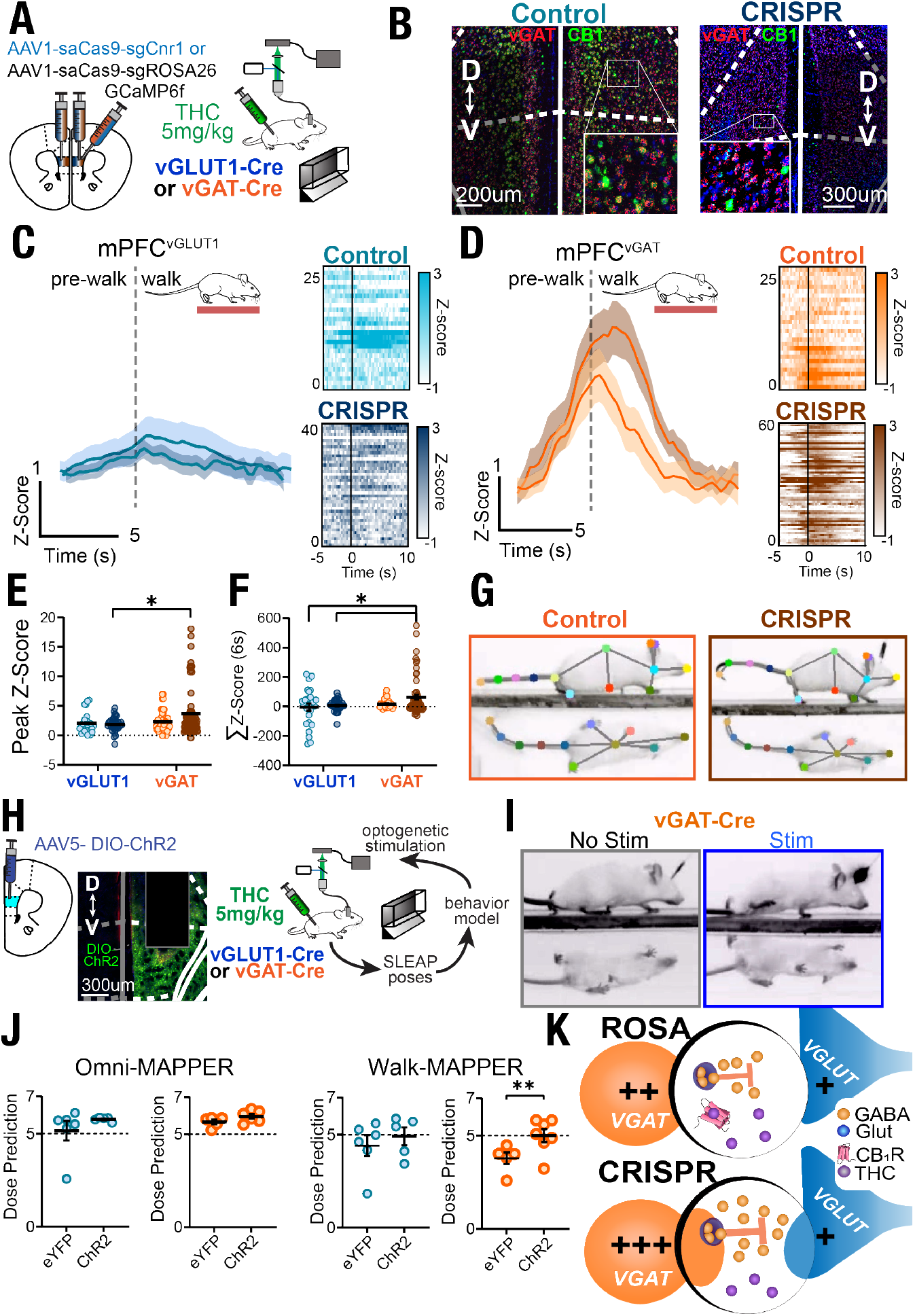
Behavioral Decoding Coupled to Closed-loop Loop Optogenetics Establishes mPFCvGAT neurons as sufficient for eliciting THC-induced disrupted behavioral kinematics: **A)** Schematic of bi-lateral AAV1-SaCas9-sgCnr1 or AAV1-SaCas9-sgROSA control injection and unilateral DIO-GCaMP6f injection with fiber implantation for fiber photometry. **B)** Microscopic images of mPFC^vGLUT1-ROSA^ and mPFCv^GLUT1-CRISPR^ tissue slices from RNAscope *in situ* hybridization. **C-D)** Fiber photometry signal at walk initiation of mPFC^vGLUT1^ (**C**) and mPFC^vGAT^ (B) ROSA control (blue and orange) and CRISPR-expressing (purple) mice, shown as calculated Z-score. **E-F)** Peak Z-score (**E**) and Σ Z-score (6 s) (**F**) over neural recordings of walk events in mPFC^vGAT-ROSA^ (orange, “Control”), and mPFC^vGAT-CRISPR^ (orange and purple), mPFC^vGLUT1-ROSA^ (blue, “Control”), and mPFC^vGLUT1-CRISPR^ (blue and purple) recordings. **G)** Image of a mPFC^vGAT-ROSA^ and mPFC^vGAT-CRISPR^ mouse walking, mid-stride, with pose estimated labels. **H)** Schematic for closed-loop optogenetic experimentation within the linear track chamber where mice treated with THC (5 mg/kg) received photo-stimulation at walk initiation. **I)** Image of a mPFC^vGAT-ChR2^ mouse walk behavior with and without stimulation within video, midstride. **J)** Comprehensive and Kinematic dose prediction of behavior from mPFC^vGLUT1-eYFP^, mPFC^vGLUT1-ChR2^, mPFC^vGAT-eYFP^, and mPFC^vGAT-ChR2^ animals. Kinematic dose prediction only applied to walk events where stimulation occurred. **K)** Cartoon Schematic of mPFC after THC administration. N=5-10, Two-Way ANOVA and Student’s unpaired T-tests, (^*^p<0.05, ^**^p<0.01).

The efficacy of this CRISPR approach was measured by using fluorescence-assisted cell sorting (FACS) of GFP-positive nuclei, labeled through a KASH peptide-GFP fusion construct targeted to the nuclear envelope, and indicated a significant knockout via insertion/deletion modifications of the *Cnr1* gene (Supplemental Figure S7). The functionality of this CRISPR CB_1_R knockout approach was further validated by using post-hoc *in situ* hybridization RNAscope analysis in each mouse (Figure 4B-C and Supplemental Figure S7). CRISPR-deleting CB_1_R from mPFC^vGAT^ neurons increased their neural activity that was time-locked to the initiation of walk behavior in THC treated mice (5 mg/kg), whereas CRISPR-deleting CB_1_R from mPFCvGLUT1 neurons did not influence their time-locked changes in activity (Figure 4D-E). As expected, the quantification of neural traces from mPFC^vGLUT1-CRISPR^ and mPFC^vGAT-CRISPR^ (peak Z-scores and Σ Z-scores (6 s)) compared to ROSA controls revealed only a minor CRISPR-dependent change in neural signal driven from the mPFC^vGAT-CRISPR^ neurons (Figure 4F-H). Conditional CRISPR knockout of *Cnr1* in mPFC GABAergic neurons increased GABAergic activity during walk events and blunted THC-induced shifts in local E/I balance, identifying CB_1_R-expressing GABAergic inhibitory neurons as mediators of THC’s circuit-level effects.

### Behavioral Decoding Coupled to Closed-loop Loop Optogenetics Establishes mPFC^vGAT^ neurons as sufficient for eliciting THC-induced disrupted behavioral kinematics

To mechanistically evaluate the involvement of mPFC glutamatergic and GABAergic neurons in this THC-induced transient activity and impairment of walking behavior, we sought to control neuronal activity during discrete locomotor events within the linear track system using an optogenetic approach. Specifically, mPFC^vGLUT1^ and mPFC^vGAT^ neurons were targeted with uni-lateral injections to express either the light-sensitive opsin Channel Rhodopsin (AAV5-EF1a-DIO-ChR2-eYFP) or an eYFP control (AAV5-EF1a-DIO-eYFP) and then implanted with an optic fiber 4 weeks prior to behavioral testing. A challenge when investigating naturalistic behavioral events is to predict the timing of their changes between distinct events as means to accurately time the optogenetic interventions. To resolve this, we constructed a closedloop optogenetic system coupled to the real time decoding of behavioral events within our linear track chamber (Figure 4I-J). Here we constructed a second supervised classification algorithm trained solely to identify walking as means to predict each “pre-walk” event that occur 30 frames immediately prior to walk initiation (*>* 95% precision and *>* 85% recall of model accuracy, Supplemental Figure S1). Next, we set the photo-stimulation parameters to trigger ramping photostimulation when a pre-walk event was identified. Photo-stimulation was triggered in a closed-loop manner using a 0.5 s ramp to 5 Hz where it remains for the duration of the walk event (5 ms pulse width and 10 s inter-trial-interval [**ITI**]) (38). Figure 4K shows that the photo-stimulation of THC-treated mPFC^vGAT-ChR2^ mice increased the predicted dose of THC calculated by the kinematic model, thus enhancing locomotor impairment compared to mPFC^vGAT-eYFP^ controls. By contrast, photo-stimulation of mPFC^vGLUT1-ChR2^ mice did not change in predicted THC-induced change in behavior by either model as compared to mPFC^vGLUT1-eYFP^ controls. The enhanced impairment of walking kinematics observed following activation of CB_1_R-expressing mPFC^vGAT-ChR2^ neurons by THC highlights the critical role of inhibitory mPFC circuits in mediating THC impairment of locomotion suggesting eCB signaling is involved (Figure 4L).

### THC potentiates the transient increase in 2-AG release by mPFC^vGLUT1^ neurons at walk initiation

THC exerts its psychoactive effects by partially activating CB_1_R, which disrupts the eCB signaling system (39). 2-AG, the most abundant eCB in the brain, is produced by several neuronal subtypes and acts in a retrograde manner to activate pre-synaptic CB_1_R as a full agonist (40, 41). To identify whether the cellular source of 2-AG production in the mPFC is linked to THC-induced impairment of locomotion, expressed the eCB sensor, GRAB_eCB2.0_, in mPFC neurons (mPFC^GRABeCB^) driven by the pan-neuronal promoter synapsin (AAV1-hSyn-GRAB-eCB2.0) (Figure 5A). mPFC^GRABeCB^ mice were treated with either vehicle or THC (1, 5, and 10 mg/kg, *i*.*p*.) 1 h before recording 2-AG dynamics during naturalistic monitoring within the linear track. As expected, LOB and crouching behavior, respectively were increased and decreased in a dose-dependent manner by

**Figure 5.**
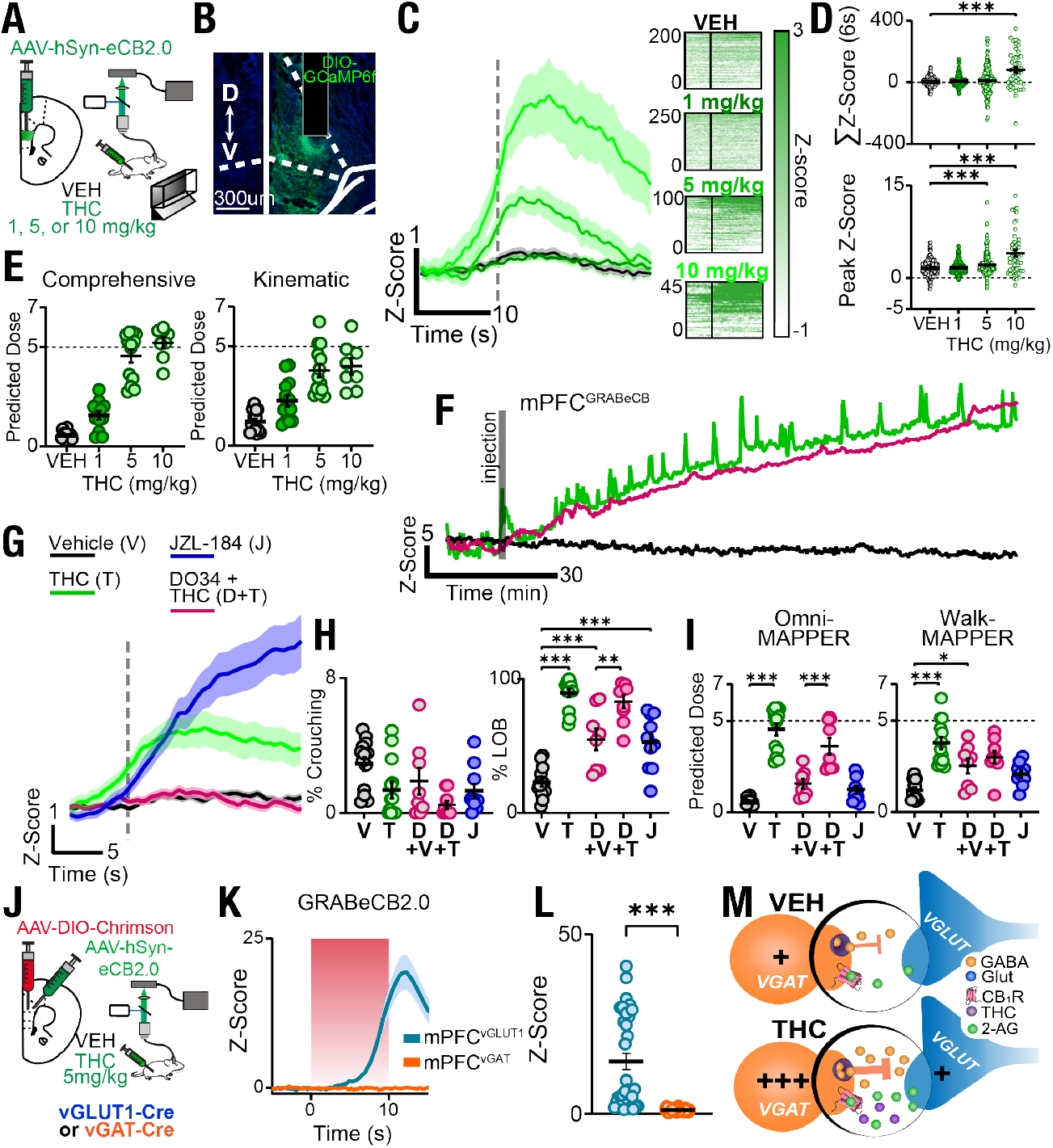
THC induces potentiation of 2-AG transients and release by mPFC^vGLUT1^ neurons at walk initiation: **A)** Schematic for GRABeCB2.0 expression and histology with optic fiber placement. **B)** Percent laying on belly and Crouching behavioral presentation across mPFC^GRABeCB^ animals treated with vehicle, 1, 5, or 10 mg/kg. **C)** Fiber photometry signal of mPFC^GRABeCB^ signal in WT mice treated with vehicle or a range of THC doses (1, 5, or 10 mg/kg) at walk initiation with their subsequent heatmaps for each walk event. **D)** Z-score (6 s) and peak Z-score from walk events. **E)** Dose prediction with Comprehensive and Kinematic dose prediction models for vehicle and THC-treated animals. **F)** Example fiber photometry trace of a 100 min experimental session with 10 mg/kg THC (green) injected systemically at 10 min. Animal was pre-treated 1 h before with 30 mg/kg DO34 before systemic injection of vehicle (black) or 5 mg/kg THC (purple). **G)** Fiber photometry signal at walk initiation after treatment with vehicle (black), 5 mg/kg THC (green), 30 mg/kg DO34 then 5 mg/kg THC (purple), or 20 mg/kg JZL-184 (maroon). **H)** Percent LOB and Crouching behavioral presentation across all eCB pharmacological treatments of vehicle (V), 10 mg/kg THC (T), 30 mg/kg DO34 + vehicle (D+V), 30 mg/kg DO34 + 10 mg/kg THC (D+T), and 20 mg/kg JZL-184 (J). **I)** Dose prediction of all eCB pharmacological treatments with total and kinematic behavioral models. **J)** Schematic for Cre-dependent expression of red-shifted opsin, ChrimsonR, and GRAB^eCB2.0^ with hSyn promoter in vGLUT1-Cre (mPFC^vGLUT1-GRABeCB^) and vGAT-Cre (mPFC^vGAT-GRABeCB^) mice. ChrimsonR stimulation at 20 Hz (5 ms pulse width, 90 s ITI). **K-L)** mPFC^vGLUT1-GRABeCB^ and mPFC^vGAT-GRABeCB^ signal following a 10 s, 20Hz stimulation (5 ms pulse width). **M)** Cartoon schematic of local mPFC circuitry during walk behavior of THC-treated mice. N=7-16 Student’s unpaired T-tests, Two-Way ANOVA with multiple comparisons (^**^p<0.01, ^***^p<0.001).

THC (Figure 5B). Similarly to mPFC^vGLUT1-GCaMP6f^ and mPFC^vGAT-GCaMP6fg^ neurons, THC induced transient production of eCBs at the initiation of the walking events with the highest activity apparent at 5 and 10 mg/kg of THC injection (Figure 5C). The integrated Z-score (6 s) and peak Z-score quantification across tested doses revealed significant dose-dependent effects at 5 and 10 mg/kg as compared to vehicle groups (Figure 5D). Predicted dosages for mPFC^GRABeCB^ animals were consistent across both models (Figure 5E). These results isolate a previously uncharacterized mechanistic link between THC-induced impairment of walking behavior and time-locked potentiation of eCB release.

To pharmacologically isolate the involvement of 2-AG in these THC-induced transient increases in mPFC^GRABeCB^ signal, we pre-treated mice with DO34 (30 mg/kg, 1 h prior), an inhibitor of the 2-AG synthesizing enzyme diacylglycerol lipase (**DAGL**). DO34 blocked the THC-induced potentiation of mPFC^GRABeCB^ signal time-locked to the initiation of walk events (Figure 5F). To further determine the involvement of 2-AG, we treated mice with JZL-184 (10 mg/kg), an inhibitor of 2-AG metabolizing enzyme, monoacylglycerol lipase (MAGL), to further amplify local increases in 2-AG levels. JZL-184 significantly prolonged the increases in mPFC^GRABeCB^ signal measured in THC-treated mice during walk events (Figure 5G). We measured the effects of either increasing or reducing 2-AG signaling by using the MAGL inhibitor, JZL-184, and the DAGL inhibitor, DO34, respectively (42, 43). Similarly to THC treatment, JZL-184 treatment increased the time spent LOB without affecting the time spent “crouching”, results that are consistent with previous findings showing that JZL-184 reduces overall movement (43, 44) (Figure 5H). DO34 also increased time spent LOB without affecting the time spent “crouching”, and its combined treatment with THC did not impact the THC response, confirming the involvement of DAGL in the THC induced transient increases in 2-AG release (45) (Figure 5H). DO34 treatment has been shown to induce mild hypolocomotion and no effects depending on readouts (44, 46, 47), results that emphasize the high sensitivity of our unbiased experimental approach to measure naturalistic mouse behaviors in a linear track. Omni-MAPPER dose prediction revealed no change from vehicle after DO34 or JZL treatment, supporting the interpretation that THC’s broad cannabimimetic effects are not driven by altered 2-AG signaling. Figure 5I shows that DO34 or JZL treatments did not affect Omni-MAPPER dose prediction results, further supporting the interpretation that THC’s broad cannabimimetic effects are not driven by altered 2-AG signaling as opposed to THC’s effect on the naturalistic behaviors reported above. Accordingly, Walk-MAPPER dose prediction increased in mice treated with DO34 treatment as single agent, further suggesting that 2-AG release in the PFC contributes specifically to discrete locomotor impairments rather than to generalized behavioral disruption (Figure 5I). Thus, our experimental approach detects changes in nuanced naturalistic mouse behavior that are distinct from canonical THC-induced patterns monitored by the cannabinoid-induced behavioral tetrad (23, 48). Mechanistically, our results show that the THC-induced potentiation of 2-AG release in the mPFC is driving its impairment effects of behavioral impairment.

Considering that concomitant and transient increases in mPFC GABAergic activity and 2-AG production is involved in the THC-induced impairment of locomotor activity, we sought to identify the cellular source of 2-AG production. Thus, we photo-stimulated either mPFC^vGLUT1^ or mPFC^vGAT^ neurons and recorded changes in GRAB_eCB2.0_ signal within the mPFC. We unilaterally co-injected viruses encoding for red-shifted opsin ChrimsonR (AAV5-FLEX-ChrimsonR-td-Tomato) and for GRAB_eCB2.0_ in the mPFC of either vGLUT1-Cre (mPFC^vGLUT1-GRABeCB^) or vGAT-Cre (mPFC^vGAT-GRABeCB^) mice and implanted an optic fiber above the region (Figure 5J). Figure 5K-L shows that photo-stimulation of mPFC^vGLUT1-GRABeCB^ neurons increased the GRAB_eCB2.0_ signal, whereas photo-stimulation of mPFC^vGAT-GRABeCB^ neurons did not. Together, our results suggest a molecular mechanism whereby 2-AG transient production from glutamatergic neurons within mPFC results in increasing the activity of CB_1_R-expressing GABAergic neurons. Our results show that THC potentiates this signaling mechanism in the mPFC inducing a robust shift in E/I balance characterized by enhanced GABA activity. These close-loop, time-locked experiments indicate that enhanced GABAergic activity is sufficient to induce disrupted kinematic parameters of mice and to impair locomotion (Figure 5M).

## 3 Discussion

Here we developed several AI analysis models that reliably identify disruption of naturalistic behaviors in mice. The effect of THC on kinematic parameters that impair locomotion is mediated by the transient potentiation of 2-AG release from glutamatergic neurons. These 2-AG transients act on CB_1_R expressed in GABAergic neurons to mediate their activity and alter the excitatory/inhibitory balance in the mPFC. This mechanism was confirmed using local THC injections in the mPFC, real-time in vivo biosensor recordings, cell type–specific CRISPR deletions and closed-loop photo-stimulation of GABAergic neurons during locomotion. Together, these results reveal a precise, cell-type-specific pathway by which THC hijacks eCB signaling to disrupt natural behavior, offering a new framework to predict and decode the subtle effects of cannabinoid agonists. Furthermore, decoding THC-induced impairment sets the groundwork for the use of unbiased pipelines for screening the bioactivity of psychoactive compounds by identifying their discrete effects on mouse behavioral signatures, which will serve as a unique preclinical experimental approach to enhance translational research. We report the natural behavior of mice in a linear track and reliably quantified the time spent in select behaviors by using both a supervised and unsupervised analysis, performing previously unappreciated natural behaviors, including “crouching” and “scanning”. Next, from these, we built Omni-MAPPER to predict the dose of THC that directly impacts these behaviors. The dose-prediction analyses presented here further highlights the potential of AI-guided behavioral analysis to revolutionize detection and interpretation of unbiased alterations in naturalistic behaviors induced by pharmacological treatment (49, 50). Leveraging random forest nonlinear regression to enhance discrimination of predicted doses, dose prediction algorithms that we report here provide an unbiased holistic approach toward deciphering drug effects on animal behavior (51).

Products containing THC are increasingly available across the USA and many additional countries (1). A better understanding of THC’s “impairing” effect has been an increasing issue for medical and legal concerns related to safety under the influence of product that contain cannabinoid agonists (1). Furthermore, some studies suggest that such products have led to motor vehicle accidents without a clear understanding of how to identify THC-induced impairment in human users (1, 52–54). Using a standardized method to detect biosignatures of THC-induced impairment in animals brings us a critical step closer to developing evidence-based tools for identifying functional impairment in humans, bridging a gap between preclinical research with real-world clinical and legal implications.

One of the most surprising results that we discovered is time-locked of 2-AG release during specific behaviors, at walk initiation, that is dose-dependently potentiated by THC treatment. Fluorescent biosensors for single ions, such as calcium, have been developed and extensively used to study the dynamic changes of such signaling molecules, only few sensors have been developed to monitor lipid signaling molecules. The recent development of fluorescence sensors based on G protein-coupled receptors that are activated by lipid mediators, here the GRABeCB2.0 has enhanced our ability to detect the time-dependent and subcellular dynamic changes of this class of hydrophobic neuromodulators. Previous studies detected changes in eCB levels *in vivo* were limited by the time dependent sampling of brain tissue using either microdialysis or tissue harvesting followed by mass spectrometry (55). Using such an approach, the behavior and drug stimulated production of eCB in select brain areas was resolved at a time resolution of minutes (56, 57). Our study provides that the first example of transient, cell-type specific, production of 2-AG that is induced by a select behavior and by closed-loop optogenetic stimulation of a neuronal subtype in a specific brain nuclei. Isolating these local circuit effects provides understanding of THC’s neuropharmacological impact on disrupting spontaneous locomotion behavior, walk-initiation, and impairment of fine motor kinematics.

The bioactivity of CB_1_R agonists in mice is traditionally measured by using a “Tetrad” set of behavioral readouts, including: hypolocomotion, analgesia, reduction in body temperature, and catalepsy, alongside paradigms that measure motivated, stress, and anxiety behaviors (58, 59). The potential therapeutic value of targeting CB_1_R and eCB signaling is only established by understanding the optimal ratio of its therapeutic versus the impact of side effects, including CB_1_R agonist induced impairment behavior.

While our study provides detailed insight into THC-induced behavioral and circuit-level effects, several limitations remain. First, our analyses were restricted to a linear track and selected natural behaviors, which capture only a subset of the wide repertoire of THC’s impact across more complex or socially relevant contexts. Second, although we resolved transient 2-AG, glutamatergic, and GABAergic signaling at high temporal resolution, other neurotransmitters and neuromodulators likely contribute to these behaviors but were not measured. Future studies should extend these approaches to more ethologically rich environments, integrate multi-region neural recordings, and incorporate additional eCB and neurotransmitter sensors to decipher any upstream or downstream neural signaling dysregulation caused by THC. Finally, our findings confidently point to a THC-dependent disinhibition effect in the mPFC, wherein enhanced 2-AG release from glutamatergic neurons inhibits select GABAergic populations. Based on prior work, vasoactive intestinal peptide- and parvalbumin-expressing interneurons are strong candidate intermediate interneurons, as these subtypes are known to regulate other local cortical GABAergic neurons (60, 61). Pinpointing the precise interneuron populations involved, and the role of CB_1_R in each will be essential for understanding how THC reshapes mPFC micro-circuit dynamics and could reveal cellular targets that mediate cannabinoid-induced behavioral impairment.

Here we show that THC fundamentally reshapes naturalistic behavior by hijacking eCB signaling, triggering transient 2-AG release from mPFC glutamatergic neurons. This signaling results in transient increases in GABAergic activity within the mPFC, likely through modulation of disinhibitory interneuron activity, disrupting locomotor initiation and altering fine motor control. Beyond these circuit-level insights, AI-guided behavioral decoding reveals subtle, previously unappreciated disruptions in naturalistic behaviors, such as crouching and scanning, that serve as uniquely sensitive biomarkers of THC exposure. Together, these findings provide a unified framework linking molecular, circuit, and behavioral effects of cannabinoids, highlighting both a mechanistic basis of impairment with a scalable, translational approach for evaluating psychoactive compounds.

## 4 Methods

### Animals

Adult (16-35g, 12-52 weeks) male and female WT, vGATCre, vGLUT1-Cre, SST-Cre, and VIP-Cre mice were group house, given ad libitum access to food and water, and maintained on a 12 hour light dark cycle (lights off at 9:00 AM, lights on at 9:00 PM). Mice were bred in a barrier facility and transferred to the holding facility at least 1 week before the start of surgical interventions or experiments. All animals were drug and test naïve, unless otherwise noted. Unless otherwise noted, statistical comparisons did not detect any sex differences, and therefore male and female mice were combined for the data presented in this manuscript. All animals were monitored daily by either facility staff or experimenter throughout the duration of the study. All procedures were approved by the Animal Care and Use Committee at the University of Washington and conformed to US National Institutes of Health guidelines. Animal studies followed the guidelines established by the Association for Assessment and Accreditation of Laboratory Animal Care (AAALAC) and were approved by the Institutional Animal Care and Use Committee (IACUC) of the University of Washington. Investigators were not blinded to experimental exposure conditions throughout assays due to the noticeable behavioral effects measured in response to THC. Animal procedures were approved by the IACUC of the University of Washington and conform to the guidelines on the care and use of animals by the National Institutes of Health.

### Pharmacological agent handling and treatments

All pharmacological treatments were performed via intraperitoneal (*i*.*p*.) route of administration. Animals received vehicle, THC (0.1, 0.3, 1, 3, 5, 10, or 30 mg/kg), SR141716 (SR1, 1 mg/kg), DO34 (30 m/kg), or JZL-184 (10 mg/kg). THC and SR141716 (SR1) were provided by the National Institute of Drugs Abuse Drug Supply Program (Bethesda, MD). DO34 and JZL-184 were purchased via Sigma-Aldrich. For injections, THC, SR1, and DO34 were dissolved in 95% ethanol and then vortexed thoroughly with equal volume cremophor and finally dissolved in sterile saline to reach a final 1:1:18 solution consisting of ethanol:cremophor:saline. JZL-184, due to its low solubility was dissolved in a 1:1:8 solution (DMSO:Cremophor:saline) with JZL-184 being dissolved in DMSO first. Appropriate controls were used for JZL-184.

### Hand-labeled behavioral tracking

Behavioral datasets and time point markings for walk behaviors in chambers other than the linear track were performed by hand. All walk events were filtered for 20 second intervals from the previous walk event to be counted as a new event. Hand-labeled behaviors were performed using the open-source software BORIS to streamline high-throughput labeling (62).

### Linear track behavioral chamber

A linear track behavioral corridor was constructed for the behavioral recording of animals from two perspectives (bottomup and side profiles) simultaneously with a single camera. This chamber was designed by Machado et al 2015 and recording was initially completed with a Basler ace camera (acA800-510um) (27). Lighting for the linear track chamber was done with 4 IR flood lights and 2 supportive IR lights positioned around the chamber. All videos were recorded at 100 frames per second using python script. Behavioral experiments that did not coincide with intracranial interventions (fiber photometry, cannulation, or optogenetics) were recorded for 15 min in the linear track. The linear track was cleaned with Clidox after every animal was exposed to the chamber and all cleaned with Clidox and then ethanol between every cage. All recordings were completed in the dark with red lights that did not obstruct the IR recordings of the camera.

### Machine learning behavioral tracking

Behavioral tracking was performed using the SLEAP tracking algorithm, a convolutional neural network published in Pereira et al 2022 (21). A specialized model was built for linear track behavior with single animal exposure. To develop the model, a subset of 1767 frames from 313 videos were selected to account for a range of sexes (51% male, 49% female), treatment conditions (Vehicle and a dose range of THC), genotypes (WT, vGAT-Cre, vGLUT1-Cre), ages (6 weeks to 6 months), and modified lighting conditions to ensure versatility of the model. Of the 1767 frames, 50% were randomly selected by SLEAP and 50% were hand-picked for difficult/valuable frames to improve model accuracy. Final model precision and recall was 97.15% and 98.74%, respectively (Supplemental Figure S1). All data analyzed were tracked with this model (recorded at 100 frames per second) and exported as .h5 files for further analysis. An analysis package suite was used to export the positional data and calculate features for respective classification methods described below.

### Supervised Classification

Identification of core behavioral responses within the linear track was performed with an automated supervised ML algorithm to identify frame-by-frame instances of walking, rearing, grooming, and laying on belly. This model was trained on 1.4 million hand-scored frames of naturalistic behavior within the linear track from a dataset of 45 videos consisting of adult (6 week to 6 month) 51% female and 49% male mice. Videos of animals were treated with either vehicle or some drug to expand the model versatility across different behavioral modifications. Model was trained on 1.85 million frames and separated test set revealed a final precision and recall of 98.3% and 97.7%, respectively. Further code was used to isolate behavioral event timestamps from the supervised model to calculate kinematic features or align neural data when applicable.

### Kinematic analysis

Analysis of natural behavioral walking kinematics within the linear track was done first by isolating natural walk events across videos with the supervised classification algorithm described above. Walk events were then isolated from the video for batch analysis. Estimated pose points of interest were used to calculate chosen metrics of locomotor kinematics commonly analyzed in previous literature. Metrics were averaged over the course of given walk events and then averaged across all walk events for a given animal. At higher dose THC treatments there were markedly less walk events which did contribute to a higher variability in addition to modified metrics. Due to concerns of multiplicity significance, kinematic features were analyzed for visualization purposes but only gait widths and dose prediction modeling were used for biological conclusions.

Isolating steps: estimated POIs from the bottom view for the right and left forepaws and hindpaws (4 total) were isolated to identify steps within the walk event. To do this, velocities were measured at every frame and plotted to reveal a wave form-like rate as paws enter and exit a stance phase. Delineation of a “step” within a walk was determined for each paw individually to be within the most recent peak speed (middle of final stride for a given walk, Supplemental Figure S2). Steps separations were used for some kinematic metrics such as stride length and step number. For kinematics features specific to a given paw, if otherwise not specified to a given paw, metrics were averaged across all four paws.

### Dose Prediction Modeling

Identification of a “presented dose” of behavior or neural signal was determined with a random forest regression algorithm. This model type was selected to account for nonlinear distributions in effects caused by varying THC dose and allow for the identification of fractional shifts in presented dose. Other regression models such as XGBoost were used and found to be less accurate than random forest regressors. All models were trained on a subset of all WT behavioral C57/bl6 videos recorded in the linear track in which animals were treated with a range of THC 0.1-7 mg/kg. Higher doses (10 and 30 mg/kg) and vehicle videos were removed as it improved the accuracy of the models. All videos for training were 15 min long and retained at that size for consistency and to account for variability in the first few min where activity was heightened. All models were trained with a 70-30 split of training to testing videos split evenly across doses where 70% of videos were used to train the algorithm and 30% were used to test the accuracy. Accuracy measurements for each model were assessed by mean square error (**MSE**).

For all models, feature importance was assessed using Scikit learn permutation importance and using a permutation analysis to assess individual contributions to the model’s accuracy (as measured by MSE). Features that contributed to the bottom 25% were removed and the model retrained to assess if model accuracy improved. If it did, the feature was discarded, otherwise it remained for the final model training. All features listed in the following Dose Prediction sub headers are the final selected features found to contribute to model accuracy.

When analyzing a novel video to assess presented dose with a dose prediction model, features for the given video were calculated in respect to the model of interest. Then, the regressor was applied to predict on the calculated features. High doses over 7 mg/kg were excluded from model training as they reduced the accuracy of the model, likely due to significant inactivity and lower transition events.

For all behavioral classification and unsupervised analyses of nuanced behaviors, review by eye ensured “humanin-the-loop” validation to ensure the model was not misidentifying specific behaviors.

### Dose Prediction Modeling of General Behaviors

Identification of general naturalistic behavioral signatures to identify THC “presented dose” for each animal was done by utilizing metrics from the previously described supervised algorithm. Core behaviors (walking, rearing, grooming, laying on belly, and other behaviors) were calculated for all training set videos at each frame and smoothed following description above. Then, the following features were calculated on each training video before being used to train a random forest regressor. The final features utilized were as follows: 1) average length of another bout, 2) average length of a walk bout, 3) average length of a rear bout, 4) average length of a groom bout, 5) average length of a lay on belly bout, 6) standard deviation of length of another bout, 7) standard deviation of length of a walk bout, 8) standard deviation of length of a rear bout, 9) standard deviation of length of a groom bout, 10) standard deviation of length of a lay on belly bout, 11) number of other bouts, 12) number of walk bouts, 13) number of rear bouts, 14) number of groom bouts, 15) number of lay on belly bouts, 16) percent time spent in other behavior, 17) percent time spent in walk behavior, 18) percent time spent in rear behavior, 19) percent time spent in groom behavior, 20) percent time spent in lay on belly behavior, and 21) total number of behavioral transitions.

### Dose Prediction Modeling of Kinematic Behaviors

First, walk events across the training dataset were identified using the Supervised algorithm to identify naturalistic walk events. Then, kinematic features were calculated following the description outlined in Kinematic Analysis. All features were calculated solely during walk events and therefore provided a higher within-video confidence at lower dose THC treatments where the animals performed a greater number of walk events. The final features utilized were as follows: 1) back height, 2) belly height, 3) nose height, 4) tail base height, 5) tail tip height, 6) average tail base yaw, 7) average head yaw, 8) average body yaw, 9) average right forepaw yaw, 10) average left forepaw yaw, 11) average right hind paw yaw, 12) average left hind paw yaw, 13) average hind paw gait width, 14) average forepaw gait width, 15) average body angle, 16) average body pitch, 17) average head pitch, 18) average walk bout distance, 19) average walk bout speed, 20) average number of steps, 21) average stride speed, 22) average stride distance, 23) average stride time, 24) average paw speed, 25) maximum walk speed, 26) minimum walk speed, 27) standard deviation of bout distance, 28) standard deviation of bout speed, 29) standard deviation of number of steps, 30) standard deviation of stride speed, 31) standard deviation of stride distance, 32) standard deviation of stride time, 33) standard deviation of right hind paw yaw, 34) standard deviation of left hind paw yaw, 35) standard deviation of right forepaw yaw, 36) standard deviation of left forepaw yaw, 37) standard deviation of minimum speed, 38) standard deviation of maximum speed, 39) standard deviation of paw speed, 40) standard deviation of head yaw, 41) standard deviation of tail base yaw, 42) standard deviation of body yaw, 43) standard deviation of forepaw gait width, 44) standard deviation of hind paw width, 45) standard deviation of body angle, 46) standard deviation of head pitch, and 47) standard deviation of body pitch. Of the 60 original features selected, 13 were removed after being found to reduce model accuracy.

### A Unsupervised Classification

Assessment of nuanced naturalistic behaviors was performed on “other” behaviors as identified by our supervised algorithm. These behavioral frames represent all actions performed by the mice that could not be described as walking, rearing, grooming, or laying on belly by the supervised algorithm. These frames across all WT treatments, including vehicle and THC doses 0.1 mg/kg to 30 mg/kg, were isolated and 29 chosen features were selected to represent the animal’s posture in each frame. Certain features were selected to account for temporal dynamics between frames (e.g. velocity and acceleration of POIs). These 29 features, across all videos, were then used in a high dimensional clustering Kmeans algorithm to identify 12 unique clusters of behaviors. The cluster number was iterated on to determine the optimal cluster number by building Kmeans models at varying cluster numbers and recording the downstream accuracy of a dose prediction model (see below).

Within each video, cluster number, frequency and bout length were quantified relative to the total number of “other” frames within each respective video to investigate the dose-dependent effects as displayed in Figure 2. A random selection of cluster frames was selected across videos and visualized for a “human-in-the-loop” assessment of model performance and for behavioral identification. These clusters are represented in Supplemental Data where names have been given to specific clusters such as “sitting”.

### Dose Prediction Modeling of Unsupervised Behaviors

Unsupervised clusters of nuanced naturalistic behaviors revealed dose-dependent behavioral expressions that were utilized to construct a dose prediction model. The final features utilized were as follows: 1) Percentage of cluster 1 frames, 2) Percentage of cluster 2 frames, 3) Percentage of cluster 3 frames, 4) Percentage of cluster 4 frames, 5) Percentage of cluster 5 frames, 6) Percentage of cluster 6 frames, 7) Percentage of cluster 7 frames, 8) Percentage of cluster 8 frames, 9) Percentage of cluster 9 frames, 10) Percentage of cluster 10 frames, 11) Percentage of cluster 11 frames, 12) Percentage of Crouching frames, 13) Mean bout length of cluster 1, 14) Mean bout length of cluster 2, 15) Mean bout length of cluster 3, 16) Mean bout length of cluster 4, 17) Mean bout length of cluster 5, 18) Mean bout length of cluster 6, 19) Mean bout length of cluster 7, 20) Mean bout length of cluster 8, 21) Mean bout length of cluster 9, 22) Mean bout length of cluster 10, 23) Mean bout length of cluster 11, 24) Mean bout length of Crouching, 25) Standard Deviation of cluster 1 bout length, 26) Standard Deviation of cluster 2 bout length, 27) Standard Deviation of cluster 3 bout length, 28) Standard Deviation of cluster 4 bout length, 29) Standard Deviation of cluster 5 bout length, 30) Standard Deviation of cluster 6 bout length, 31) Standard Deviation of cluster 7 bout length, 32) Standard Deviation of cluster 8 bout length, 33) Standard Deviation of cluster 9 bout length, 34) Standard Deviation of cluster 10 bout length, 35) Standard Deviation of cluster 11 bout length, 36) Standard Deviation of Crouching bout length.

### Combination of Dose Prediction models

Features from the training behavioral set were calculated for all behavioral feature sets as described for the general, kinematic, and unsupervised dose prediction models. For the combined general and kinematic model in Figure 1, the features used to train the general and kinematic models were combined into a large array. This combined feature array across doses was used to train the combined general-kinematic behavioral model. Similarly, the features were all combined into one large feature set before being used to train a new random forest algorithm to predict dose for Comprehensive presented dose.

### Tissue processing

For all fiber photometry experiments, mice were transcardially perfused with a 4% paraformaldehyde solution in phosphate buffered saline. Heads were subsequently removed and were immediately placed into a same solution bath to set for 24 hours. After this time, fiberoptic implants were removed along with the remainder of the skull, and brains were placed into a cryoprotectant 30% w/v sucrose solution in PBS. Once brains had sunken to the bottom of the tubes (typically around 3-5 days), brains were mounted on cutting blocks using O.C.T. (Sakura Finetek) and sliced in 40*µ*m sections on a Leica CM1900 cryostat at -18*°*C. Slices were then placed into PBS and either immediately mounted on SuperFrost Plus slides (Fisher) for imaging or stored for up to 1 week. Mice with misplaced fiber optic implants were excluded from the study.

### Image acquisition and analysis

For histological verification of fiberoptic/ lens placement, images were obtained on a Leica DM6B epifluorescent microscope at 10x or 20x magnification. For RNAscope analysis, images were collected on an Olympus Fluoview 3000 at 20x magnification. Z-stacked images were taken with optical section thickness of 2-3*µ*M. Images were subsequently analyzed using either HALO or QuPath software. For either analysis, brain region boundaries were drawn in accordance with the Allen Brain Institute mouse reference atlas. For HALO, DAPI stained nuclei were used to mark cells, and cell boundaries were set to ∼8 *µ*M radius from nucleus center. Thresholds were set for detection in each channel and this threshold was applied to all analyzed images in a given cohort. In order to account for background noise, a neuron had to have *>* 2 transcripts in order to be classified as positive for a specific marker. For QuPath analysis, DAPI was similarly used to define neurons and cell boundaries were set to ∼8 *µ*M radius from nucleus center. Individual channels were set via the classifier module, and these classifier settings were held consistent for all images within a cohort. The classifiers were set by individually sorting through each cell automatically registered by the QuPath software and identifying the first “false positive” neuron, using this as the threshold cut off for subsequent identification of positive neurons.

### Stereotaxic surgeries

Mice were anaesthetized in an induction chamber (1-4% isoflurane) and placed into a stereotaxic frame (Kopf Instruments, model 1900) where they were mainlined at 1-1.5% isoflurane. For all viral infections, a Hamilton Neuros Syringe was used, and virus was infused at a rate of 75 nl/minute. For fiber photometry experiments, 500 nl AAV5-DIO-GCaMP6f (Addgene) or 500 nl AAV1-hSyn-GRAB_eCB2.0_ was injected into the PrL of the mPFC of vGAT-Cre, vGLUT1-Cre, VIP-Cre, SST-Cre, or WT mice, with final coordinates of A/P 1.8, D/V -2.0, M/L 0.4. After appropriate infusion of virus, 400uM fiber optic implantation was performed and bonded to the skull with metabond. For CRISPR/ photometry experiments, AAV1-FLEX-sgCNR1-SaCas9 was infused into the PrL at the same coordinates above in vGAT-Cre and vGLUT1-Cre mice. Mice were left for a minimum of 6 weeks to improve viral expression and provide time for recovery before performing a second surgery to inject AAV5-DIO-GCaMP6f and implant a 400uM fiber optic with a stainless-steel ferule (Doric). All implants were secured using Metabond (Parkell). For optogenetics experiments, 500nl of AAV5-DIO-ChR2-eYFP (Addgene) was injected into the PrL of the mPFC of vGAT-Cre, vGLUT1Cre, VIP-Cre, and SST-Cre mice. After appropriate infusion of virus, ipsilateral 200µM fiberoptics were implanted over the PrL (final coordinates of A/P +1.8, D/V -2.0, M/L + 0.4). The optic fiber was subsequently attached to the skull with Metabond. Intracranial cannulation was performed by drilling bi-lateral holes at the coordinates of A/P +1.8, D/V -2.0, M/L *±*0.4 whereby a bi-lateral cannula was carefully implanted over the PrL. The cannula was attached to the skill with metabond. All mice were given at least 4 weeks of recovery time post-surgery before experimentation.

### Fiber photometry

Fiber photometry recordings were obtained throughout the duration of all behavioral tests presented in this manuscript. Prior to recordings, a fiberoptic cable was attached to the fiberoptic implant using a plastic or ceramic ferrule sleeve (Doric, ZR2.5). For all photometry recordings, we used a Tucker-Davis Technologies RZ10x processor. A 531-Hz sinusoidal 470 nm LED light (Lx465) was used to excite GCaMP6f or GRAB_eCB2.0_ and evoke Ca^2^-dependent emission. A 211-Hz sinusoidal 405 LED (Lx405) light was used as the Ca^2^-independent isosbestic control emission. LED intensity was measured at the tip of the cable with a dummy implant attached and adjusted to 30µW before each day of recording. GCaMP6f fluorescence traveled through the same optic fiber before being bandpass filtered (525 *±* 25 nm, Doric, FMC4), transduced by a femtowatt silicon photoreceiver (Newport, 2151) and recorded by a real-time processor (TDT, RZ10). The envelopes of the 531-Hz and 211-Hz signals were extracted in realtime by the TDT program Synapse at a sampling rate of 1017.25 Hz. For ChrimsonR stimulation experiments, a 635nm laser was used with a custom filter cube (Doric) at 5-7mW of intensity to deliver 1-20hz pulsed light through the same fiberoptic cable used for photometry recordings.

### Photometry analysis

Custom MATLAB scripts were developed for analyzing fiber photometry data in context of mouse behavior and can be accessed via GitHub. The isosbestic 405nm excitation control signal was scaled to the 470nm excitation signal, then this refitted 405nm signal was subtracted from the 470 nm excitation signal to remove movement artifacts from intracellular Ca^2+^-dependent GCaMP6f or GRAB_eCB2.0_ signal. Baseline drift was evident in the signal due to slow photobleaching artifacts, particularly during the first several min of each hourlong recording session. A double exponential curve was fitted to the raw trace and subtracted to correct for baseline drift. After baseline correction, Δf/f was calculated by subtracting the median baseline subtracted signal from the baseline subtracted signal and dividing by the median raw signal. All photometry data presented at the onset of a behavioral event in this manuscript is Z-scored to a 10-30 second window prior to cue onset.

### Electrophysiology

Coronal brain slices were prepared at 250 *µ*M on a vibrating Leica VT1000S microtome using standard procedures. Mice were anesthetized with Isoflurane, and transcardially perfused with ice-cold and oxygenated cutting solution consisting of (in mM): 93 N-Methyl-D-glucamine (NMDG), 2.5 KCL, 20 HEPES, 30 NaHCO3, NaH2PO4, 10 MgSO4*·*7H20, 0.5 CaCl2 *·* 2H20, 25 glucose, 3 Na^+^-pyruvate, 5 Na^+^- ascorbate, and 5 N-acetylcysteine. Following collection of coronal sections, the brain slices were transferred to a 34*°*C chamber containing oxygenated cutting solution for a 10-minute recovery period. Slices were then transferred to a holding chamber consisting of (in mM) 92 NaCl, 2.5 KCl, 20 HEPES, 2 MgSO4·7H20, 1.2 NaH2PO4, 30NaHCO3, 2 CaCl2 · 2H20, 25 glucose, 3 Na-pyruvate, 5 Na-ascorbate, 5 N-acetylcysteine and were allowed to recover for ≥ 30 min. For recording, slices were perfused with oxygenated artificial cerebrospinal fluid (ACSF; 31-33*°*C; 300-303 milliosmols) consisting of (in mM): 113 NaCl, 2.5 KCl, 1.2 MgSO4 7H20,2.5 CaCl2 ·6H20, 1 NaH2PO4, 26 NaHCO3, 20 glucose, 3 Na^+^-pyruvate, 1 Na^+^-ascorbate, at a flow rate of 2-3ml/min. For recordings of inhibitory currents, 10 *µ*M CNQX was added to the external solution. Stocks of 50mM THC were made in ethanol and diluted 1:5000 in ACSF or HEPES for a final concentration of 10 *µ*M. 0.05% w/v Bovine Serum Albumin (Sigma-Aldrich) was added to ACSF or HEPES to help the drugs from precipitating out of solution. For all drug experiments, slices were incubated in a HEPES bath containing the drugs for >30 min before transferring over to the slice rigs for recording.

mPFC neurons were initially voltage clamped in whole-cell configuration using borosilicate glass pipettes (2-4MΩ). For recordings of excitatory currents, pipettes were filled with internal solution containing (in mM): 125 K^+^- gluconate, 4 NaCl, 10 HEPES, 4 MgATP, 0.3 Na-GTP, and 10 Na-phosphocreatine (pH 7.30-7.35). The patch pipette also included 50 *µ*M picrotoxin to block GABAA currents. For recordings of inhibitory currents, pipettes were filled with 115 CsCl2, 5 NaCl, 10 HEPES, 5 QX-314, 4 Mg-ATP, 0.3 Na-GTP, and 10 Glucose (pH 7.30-7.35). Following break-in to the cell, we waited ≥3 min to allow for exchange of internal solution and stabilization of membrane properties. Neurons with an access resistance of > 30MΩ or that exhibited greater than a 20% change in access resistance during the recording were not included in our datasets. For all voltage clamp experiments, neurons were held at -70mV.

### *Ex-vivo* Optogenetics

To assess how THC treatment modulated eCB regulation of excitatory signaling in the mPFC, VGAT-Cre mice were injected with 550 nL of AAV5-CaMKII-ChR2(H134R)-eYFP into the contralateral mPFC, and 550 nL of DIO-mCherry was injected ipsilaterally to allow us to record from either VGAT+ neurons or putative pyramidal neurons (mCherry negative). To assess how THC treatment modulated eCB regulation of inhibitory signaling in the mPFC, VGAT-Cre mice were injected with 550 nL of AAV5-DIO-ChR2(H134R)-eYFP ipsilaterally and recordings were obtained from putative pyramidal neurons (eYFP negative). 3-5 weeks of viral expression was allowed prior to sacrificing the mice. For all experiments using THC, corresponding vehicle recordings were obtained from the same mice. For optogenetic recordings of input/output curves, we used a Thorlabs LEDD1B T-Cube driver and obtained separate recordings of 470nm wavelength oEPSCs / oIPSCs at 7 output levels corresponding to 5.7, 2.7, 1.6, 1, 0.5, 0.2, and 0.1 mW of LED intensity. PPR recordings of oEPSCs / oIPSCs were obtained in voltage-clamp with an inter-stimulus interval of 50ms. PPR is reported as a ratio between the amplitude of the second oEPSC / oIPSCs divided by the first. Recordings of DSE or DSI were obtained following at 10 s voltage step to +30mV. A baseline of 10 oEPSCs / oIPSCs were taken prior to the depolarizing step, and all data is plotted as an oEPSC / oIPSCs amplitude normalized to the baseline period. A light exposure time of 2ms was used for all optogenetic experiments.

### *In situ* hybridization

For quantification of mRNA transcripts using RNAscope (ACD Bio), mice were briefly anesthetized using isoflurane and subsequently rapidly decapitated. Brains were immediately removed from the skull and were placed liquid nitrogen for rapid freezing. Then, brains were transferred to a 80*°*C freezer for long term storage. Prior to sectioning, brains were placed in the cryostat compartment for >30 min to allow them to come up to temperature. Following mounting using O.C.T., brains were slice at 16*µ*m and directly mounted onto SuperFrost Plus slides. Slides were stored at -80*°*C.

FISH was performed according to the RNAscope 2.0 Fluorescent Multiple Kit User Manual for Fresh Frozen Tissue (Advanced Cell Diagnostics) as previously described. Briefly, slides were fixed in 40% PFA and then serially dehydrated using increasing concentrations of ethanol (50%, 75%, 100% and 100%). Slices were then treated with Protease IV and allowed to incubate in a 40*°*C hybridization oven for 30 min. Following serial rinses in a wash buffer, slices were incubated with probes directed against a series of probes outlined in given experiments. Samples subsequently went through sequential amplification steps, and lastly application of Opal dyes corresponding to each channel (TSA Vivid Fluorophore 520 (PN 323271), TSA Vivid Fluorophore 570 (PN 323272) and TSA Vivid Fluorophore 650 (PN 323273)). Slides were counterstained with DAPI, and coverslips were mounted with Vectashield Hard Set mounting medium (Vector Laboratories).

### Intracranial cannulation

Male and female mice were implanted with bi-lateral guide cannulas with an internal cannula and a cap (RWD: specs) into the Pre-frontal cortex (AP: 1.8, DV: -2.0, ML: ±0.4). Cannulas were bonded to the skull with metabond and covered in a 5 mM quinine-Cremophor (50:50) solution daily to stop the animals from chewing through the material. Animals were given a 1-week recovery time before experimentation. All animals underwent a vehicle (0.5 : 0.5 : 1 : 17 = DMSO : ethanol : Cremophor : 50mg/ml BSA in saline) test day before any other treatment experiment. For all experimental days, animals were scruffed gently and held in a prone position while the head cap and dummy cannula were removed. An internal infusion cannula was then placed to infuse the desired drug at a rate of 1 uL over 2.5 min, bilaterally. Afterward, animals remained in the scruff for 1 min to allow for diffusion of the treatment before replacing with the dummy cannula and head cap. Then, mice were placed into a homecage like environment alone for 5 min prior to being placed into the linear track for a 15 min recording. Treatments given included THC at 10 ug (conc), THC at 1ug (conc), DO34, or 20 *µ*g JZL per hemisphere. Due to the high solubility of JZL-184, it had to be administered in 100% DMSO and tubing was changed between every animal.

Mice were placed into the linear track for 15 min to record a baseline for behavior and then were removed to administer the treatment (250 nl over 2.5 min) with 5 min of recovery ( ∼ 10 min) before being placed back into the linear track for 15 min. Treatment groups were counterbalanced, either receiving a vehicle or a 5 mg THC infusion. After treatment, animals were placed back in their home cage.

### Closed-loop optogenetic stimulation

To perform Closed-Loop optogenetic stimulation, two identical cameras (as described above) were setup to face the linear track chamber. One camera was used with the same computer as for all experimentation which recorded at the normal 100 fps for behavioral analysis. The other camera was connected to a second computer. This second computer was equipped with an Arduino controller and subsequent LED with a patch cable for connecting to a prepared mouse. The Arduino code was constructed to set the stimulation frequency (20 Hz), the pulse width (5 ms), the total stimulation time (3 s) and the ramp time (0.5 s) which was written to replicate the ramping in fiber photometry activity seen in Figures 3 and 4. The closed loop script that controlled the camera video capture was written in python and run to capture frames at 30 fps (the fastest possible while performing frame inference without dropping frames). The code batched frames into a sliding window of 15 frames at a time where each frame was passed through a pre-built “pre-walk” Random Forest Classification algorithm which identified behavior as either a 1 (pre-walk), 2 (walking), or 0 (not walk-related). When a threshold of 12 1’s was met within the sliding window, stimulation was triggered. This number of 12 was optimized through trial and error with the efficiency of the algorithm in real time testing. Stimulation was triggered by sending a pulse signal to the Arduino controller which then activated the LED stimulation through the patch cable to the mouse.

Male and female vGLUT1-Cre and vGAT-Cre mice were unilaterally injected with AAV-DIO-eYFP or AAV-DIO-ChR2-eYFP (600 nl) at the mPFC coordinates (AP = 1.8, DV = -2.0, ML = 0.4) before being implanted with an optic fiber. 4 weeks after injection and implantation, mice were habituated to the linear track on two days, 10 min each. On the first test day, animals’ fibers were attached to the LED patch cable and placed in the linear track. The closed loop script was then run to initialize the Arduino and LEDs while the loop code began. After a manual 15 second delay, the video recording from the original camera began recording for a total 15 min session. After 15 min, the closed loop script was halted and data was exported for further analysis. Due to any potential inaccuracy in the closedloop walk identification, all videos were checked by eye to isolate all walk events where stimulation occurred. The pose estimation data for these walk events was then separated and analyzed for the resulting kinematic dose prediction models.

### Pre-walk Random Forest Classification Algorithm

To develop the “pre-walk” classification algorithm, we used the same hand-labeled data as used for the supervised classification algorithm. Instead, we isolated only the events labeled for walking, and then all batches of 30 frames prior to each walk event as a “pre-walk”. 30 frames was the speed at which frames were assessed during live recording, equating to 1 second prior to a respective walk event. Then, we used the same train/split for our data and trained the algorithm to have a final precision and recall for pre-walk (96.4% and 97.2%) and walk (98.8% and 87.6%). Then, the algorithm was uploaded and validated with several animals in the linear track closed loop system described above.

Comprehensive behavioral dose prediction of THC 5 mg/kg treatment conditions for eYFP and ChR2 was compared to reveal that mPFC stimulation of mPFC^vGLUT1-ChR2^ or mPFC^vGAT-ChR2^ neurons did not significantly modify general THC-induced behavioral effects. Before applying the kinematic dose prediction model, all stimulated walk events were separated from those where stimulation did not occur due to either events occurring during the ITI or them being missed by the closed-loop system. Kinematic dose prediction of stimulated walk events revealed a significant increase in the predicted dose after enhancing the activity of mPFC^vGAT-ChR2^ neurons.

### ChrimsonR stimulation and GRAB^eCB2.0^ fiber photometry

Male and female vGLUT1-Cre and vGAT-Cre mice were uni-laterally injected with a viral mixture of AAV-DIO-ChrimsonR (450 nl) and AAV-hSyn-GRAB_eCB2.0_ (450 nl) at a rate of 75 nl/min at the mPFC coordinates (AP = 1.8, DV = -2.0, ML = 0.4). Then, immediately after injection, they were implanted with an optic fiber, affixed to the skull with metabond dental cement. Four weeks after injection and implantation, mice were placed in a MedPC behavioral chamber and their optic fiber attached to a TDT stimulation LED stimulation system. Over an 8 min session, animals received 565 nm wavelength stimulation at 1:00, 2:30, 4:00, 5:30, and 7:00 so as to leave 90 s in between stimulations while recording 470 and 405 nm wavelengths. Stimulation in the main figure was performed at 20 Hz frequency with 5 ms pulse width for a total of 10 seconds. Animals were pre-treated 1 hour prior to testing with either vehicle or 5 mg/kg THC and there was always at least 1 week in between tests for each animal. Fiber photometry data was analyzed as described above with the stimulation time points used as epoch references.

### Data statistical analysis and visualization

All data was plotted using either python scripts described above or by plotting exported data into GraphPad Prism. All statistical analyses that resulted in p-value calculation and all post-hoc analyses were conducted within GraphPad Prism and are described with their respective figures throughout the manuscript.

## Acknowledgments

We thank the Molecular Genetics Resource Core for the Center in Neurobiology of Addiction, Pain, and Emotion and its director, Dr. Selena Schattauer, for generating the GRABeCB2.0 and CRISPR viruses used in this manuscript. We thank Dr. Azra Suko for lab management and organization, as well as Carina Pizzano and Bailey Wells for colony management. A special thanks to John English, JD for his contributions to the physical linear track chamber construction. Lastly, we thank the entire Bruchas lab, Stella lab, and Land lab as well as other members of the NAPE center at the University of Washington for resources and critical feedback.

## Author contributions

Conceptualization: AE, MRB, NS, BBL, DJM

Methodology: AE, MRB, NS, BBL, DJM, AL, LZ

Data Analysis: AE, DJM, AL, VC, GC, IW

Investigation: AE, DJM, KY, YE, MA, AS, JPB, FU, RS

Visualization: AE, MRB, NS, BBL, DJM

Funding acquisition: AE, MRB, NS, BBL

Project administration: AE, MRB, NS, BBL

Supervision: AE, MRB, NS, BBL

Writing – original draft: AE

Writing – review & editing: AE, MRB, NS, BBL, DJM

## Data and materials availability

All data and code associated with this project will be uploaded to a public repository upon acceptance of this manuscript.

## Bibliography

1. S. D. Pennypacker, K. Cunnane, M. C. Cash, and E. A. Romero-Sandoval. Potency and therapeutic thc and cbd ratios: Us cannabis markets overshoot. Frontiers in Pharmacology, 13:921493, 2022.

2. M. Asbridge, J. A. Hayden, and J. L. Cartwright. Acute cannabis consumption and motor vehicle collision risk: systematic review of observational studies and meta-analysis. Bmj, 344:e536, 2012.

3. G. A. Van Den Elsen, L. Tobben, A. I. Ahmed, R. J. Verkes, C. Kramers, R. M. Marijnissen, et al. Effects of tetrahydrocannabinol on balance and gait in patients with dementia: A randomised controlled crossover trial. Journal of psychopharmacology, 31: 184–191, 2017.

4. N. Stella. Thc and cbd: Similarities and differences between siblings. Neuron, 111: 302–327, 2023.

5. D. McCartney, A. S. Suraev, P. T. Doohan, C. Irwin, R. C. Kevin, R. R. Grunstein, et al. Effects of cannabidiol on simulated driving and cognitive performance: a dose-ranging randomised controlled trial. Journal of psychopharmacology, 36: 1338–1349, 2022.

6. M. R. Domenici, S. C. Azad, G. Marsicano, A. Schierloh, C. T. Wotjak, H.-U. Dodt, et al. Cannabinoid receptor type 1 located on presynaptic terminals of principal neurons in the forebrain controls glutamatergic synaptic transmission. Journal of Neuroscience, 26: 5794–5799, 2006.

7. K. Monory, H. Blaudzun, F. Massa, N. Kaiser, T. Lemberger, G. Schütz, et al. Genetic dissection of behavioural and autonomic effects of δ9-tetrahydrocannabinol in mice. PLoS biology, 5:e269, 2007.

8. V. De Giacomo, S. Ruehle, B. Lutz, M. Häring, and F. Remmers. Cell type-specific genetic reconstitution of cb1 receptor subsets to assess their role in exploratory behaviour, sociability, and memory. European Journal of Neuroscience, 2020.

9. K. Rosenberg-Katz, T. Herman, Y. Jacob, N. Giladi, T. Hendler, and J. M. Hausdorff. Gray matter atrophy distinguishes between parkinson disease motor subtypes. Neurology, 80: 1476–1484, 2013.

10. R. G. Mair, M. J. Francoeur, E. M. Krell, and B. M. Gibson. Where actions meet outcomes: Medial prefrontal cortex, central thalamus, and the basal ganglia. Frontiers in Behavioral Neuroscience, 16:928610, 2022.

11. M. Laubach, M. S. Caetano, and N. S. Narayanan. Mistakes were made: neural mechanisms for the adaptive control of action initiation by the medial prefrontal cortex. Journal of Physiology-Paris, 109: 104–117, 2015.

12. G. Marsicano and B. Lutz. Expression of the cannabinoid receptor cb1 in distinct neuronal subpopulations in the adult mouse forebrain. European journal of neuroscience, 11: 4213–4225, 1999.

13. M. Tjandrasuwita, J. J. Sun, A. Kennedy, S. Chaudhuri, and Y. Yue. Interpreting expert annotation differences in animal behavior. arXiv preprint, 2021.

14. N. H. Sachuriga, Y. Takamura, J. Matsumoto, M. Ferreira Pereira de Araújo, T. Ono, and H. Nishijo. Neuronal representation of locomotion during motivated behavior in the mouse anterior cingulate cortex. Front Syst Neurosci, 15: 655110, 2021.

15. S. Hardung, Z. Jäckel, and I. Diester. Prefrontal contributions to action control in rodents. In International Review of Neurobiology, pages 373–393. Elsevier, 2021.

16. L. Wang, M. Gao, Q. Wang, L. Sun, M. Younus, S. Ma, et al. Cocaine induces locomotor sensitization through a dopamine-dependent vta-mpfc-fra cortico-cortical pathway in male mice. Nature Communications, 14:1568, 2023.

17. A. Dong, K. He, B. Dudok, J. S. Farrell, W. Guan, D. J. Liput, et al. A fluorescent sensor for spatiotemporally resolved endocannabinoid dynamics in vitro and in vivo. Technical report, 2020. bioRxiv. 2020.2010.2008.329169. bioRxiv. 2020.2010.2008.329169.

18. S. Singh, D. Sarroza, A. English, M. McGrory, A. Dong, L. Zweifel, et al. Pharmacological characterization of the endocannabinoid sensor grabecb2. 0. Cannabis and Cannabinoid Research, 9: 1250–1266, 2024.

19. A. Mathis, P. Mamidanna, K. M. Cury, T. Abe, V. N. Murthy, M. W. Mathis, et al. Deeplabcut: markerless pose estimation of user-defined body parts with deep learning. Nature neuroscience, 21: 1281–1289, 2018.

20. T. D. Pereira, D. E. Aldarondo, L. Willmore, M. Kislin, Wang Ss-H, M. Murthy, et al. Fast animal pose estimation using deep neural networks. Nature methods, 16: 117–125, 2019.

21. T. D. Pereira, N. Tabris, A. Matsliah, D. M. Turner, J. Li, S. Ravindranath, et al. Sleap: A deep learning system for multi-animal pose tracking. Nature methods, 19: 486–495, 2022.

22. C. Weinreb, J. E. Pearl, S. Lin, M. A. M. Osman, L. Zhang, S. Annapragada, et al. Keypoint-moseq: parsing behavior by linking point tracking to pose dynamics. Nature Methods, 21: 1329–1339, 2024.

23. M. Metna-Laurent, M. Mondésir, A. Grel, M. Vallée, and P. V. Piazza. Cannabinoid-induced tetrad in mice. Current Protocols in Neuroscience, 80(9): 51–59, 2017. 59. .59. 10.

24. J. L. Wiley. Sex-dependent effects of Δ9-tetrahydrocannabinol on locomotor activity in mice. Neuroscience letters, 352: 77–80, 2003.

25. C. Parks, B. C. Jones, B. M. Moore, and M. K. Mulligan. Sex and strain variation in initial sensitivity and rapid tolerance to Δ9–tetrahydrocannabinol. Cannabis and Cannabinoid Research, 5: 231–245, 2020.

26. B. R. Martin, R. L. Balster, R. K. Razdan, L. S. Harris, and W. L. Dewey. Behavioral comparisons of the stereoisomers of tetrahydrocannabinols. Life Sciences, 29: 565–574, 1981.

27. A. S. Machado, D. M. Darmohray, J. Fayad, H. G. Marques, and M. R. Carey. A quantitative framework for whole-body coordination reveals specific deficits in freely walking ataxic mice. Elife, 4:e07892, 2015.

28. A. S. Machado, H. G. Marques, D. F. Duarte, D. M. Darmohray, and M. R. Carey. Shared and specific signatures of locomotor ataxia in mutant mice. ELife, 9:e55356, 2020.

29. A. Torrens, V. Vozella, H. Huff, B. McNeil, F. Ahmed, A. Ghidini, et al. Comparative pharmacokinetics of Δ9tetrahydrocannabinol in adolescent and adult male mice. Journal of Pharmacology and Experimental Therapeutics, 374: 151–160, 2020.

30. G. Vervoort, A. Bengevoord, E. Nackaerts, E. Heremans, W. Vandenberghe, and A. Nieuwboer. Distal motor deficit contributions to postural instability and gait disorder in parkinson’s disease. Behavioural brain research, 287: 1–7, 2015.

31. C. R. Kasten, Y. Zhang, and S. L. Boehm. Acute cannabinoids produce robust anxiety-like and locomotor effects in mice, but long-term consequences are age-and sex-dependent. Frontiers in behavioral neuroscience, 13:32, 2019.

32. J. A. Marusich and J. L. Wiley. Δ9-tetrahydrocannabinol discrimination: Effects of route of administration in mice. Drug and Alcohol Dependence Reports, 9:100205, 2023.

33. A. English, F. Uittenbogaard, A. Torrens, D. Sarroza, A. Slaven, D. Piomelli, et al. A preclinical model of thc edibles that produces high-dose cannabimimetic responses. Technical report, 2022. bioRxiv. 2022.2011.2023.517743. bioRxiv. 2022.2011.2023.517743.

34. N. S. Narayanan and M. Laubach. Top-down control of motor cortex ensembles by dorsomedial prefrontal cortex. Neuron, 52: 921–931, 2006.

35. C. Risterucci, D. Terramorsi, A. Nieoullon, and M. Amalric. Excitotoxic lesions of the prelimbic-infralimbic areas of the rodent prefrontal cortex disrupt motor preparatory processes. European Journal of Neuroscience, 17: 1498–1508, 2003.

36. V. Pearson-Dennett, G. Todd, R. A. Wilcox, A. P. Vogel, J. M. White, and D. Thewlis. History of cannabis use is associated with altered gait. Drug and alcohol dependence, 178:215–222, 2017.

37. A. C. Hunker, M. E. Soden, D. Krayushkina, G. Heymann, R. Awatramani, and L. S. Zweifel. Conditional single vector crispr/sacas9 viruses for efficient mutagenesis in the adult mouse nervous system. Cell reports, 30.e4306:4303–4316, 2020.

38. E. Cela, A. R. McFarlan, A. J. Chung, T. Wang, S. Chierzi, K. K. Murai, et al. An optogenetic kindling model of neocortical epilepsy. Scientific Reports, 9:5236, 2019.

39. A. Busquets-Garcia, J. Bains, and G. Marsicano. Cb 1 receptor signaling in the brain: extracting specificity from ubiquity. Neuropsychopharmacology, 43: 4–20, 2018.

40. N. Stella, P. Schweitzer, and D. Piomelli. A second endogenous cannabinoid that modulates long-term potentiation. Nature, 388: 773–778, 1997.

41. A. Chakrabarti, J. E. Ekuta, and E. S. Onaivi. Neurobehavioral effects of anandamide and cannabinoid receptor gene expression in mice. Brain research bulletin, 45: 67–74, 1998.

42. D. Ogasawara, H. Deng, A. Viader, M. P. Baggelaar, A. Breman, H. den Dulk, et al. Rapid and profound rewiring of brain lipid signaling networks by acute diacylglycerol lipase inhibition. Proceedings of the National Academy of Sciences, 113: 26–33, 2016.

43. A. Seillier, D. D. Aguilar, and A. Giuffrida. The dual faah/magl inhibitor jzl195 has enhanced effects on endocannabinoid transmission and motor behavior in rats as compared to those of the magl inhibitor jzl184. Pharmacology Biochemistry and Behavior, 124: 153–159, 2014.

44. J. L. Wilkerson, G. Donvito, T. W. Grim, R. A. Abdullah, D. Ogasawara, B. F. Cravatt, et al. Investigation of diacylglycerol lipase alpha inhibition in the mouse lipopolysaccharide inflammatory pain model. The Journal of pharmacology and experimental therapeutics, 363: 394–401, 2017.

45. G. Bedse, N. D. Hartley, E. Neale, A. D. Gaulden, T. A. Patrick, P. J. Kingsley, et al. Functional redundancy between canonical endocannabinoid signaling systems in the modulation of anxiety. Biological psychiatry, 82: 488–499, 2017.

46. B. C. Shonesy, R. J. Bluett, T. S. Ramikie, R. Báldi, D. J. Hermanson, P. J. Kingsley, et al. Genetic disruption of 2-arachidonoylglycerol synthesis reveals a key role for endocannabinoid signaling in anxiety modulation. Cell reports, 9: 1644–1653, 2014.

47. N. Loomba, A. Cao, S. Charles, I. Kandil, M. Kwon, and S. Patel. Endocannabinoid modulation of defensive state transitions to innate and learned threat. Psychopharmacology, pages 1–14, 2025.

48. C. F. Moore and E. M. Weerts. Cannabinoid tetrad effects of oral Δ9-tetrahydrocannabinol (thc) and cannabidiol (cbd) in male and female rats: sex, dose-effects and time course evaluations. Psychopharmacology, pages 1–12, 2021.

49. Y. Voskobiynyk and J. T. Paz. Ai-nalyzing mouse behavior to combat epilepsy. Epilepsy Currents, 23: 315–317, 2023.

50. E. I. Nwokedi, R. S. Bains, L. Bidaut, S. Wells, X. Ye, and J. M. Brown. Unsupervised detection of mouse behavioural anomalies using two-stream convolutional autoencoders. arXiv preprint, 2021.

51. R. Rahman, S. R. Dhruba, S. Ghosh, and R. Pal. Functional random forest with applications in dose-response predictions. Scientific reports, 9:1628, 2019.

52. W. M. Bosker, E. Theunissen, S. Conen, K. Kuypers, W. Jeffery, H. Walls, et al. A placebo-controlled study to assess standardized field sobriety tests performance during alcohol and cannabis intoxication in heavy cannabis users and accuracy of point of collection testing devices for detecting thc in oral fluid. Psychopharmacology, 223: 439–446, 2012.

53. J. D. Brown. Potential adverse drug events with tetrahydrocannabinol (thc) due to drug–drug interactions. Journal of clinical medicine, 9:919, 2020.

54. J. D. Hinckley and C. Hopfer. Marijuana legalization in colorado: increasing potency, changing risk perceptions, and emerging public health concerns for youth. Adolescent Psychiatry, 11: 95–116, 2021.

55. A. G. Zestos and R. T. Kennedy. Microdialysis coupled with lc-ms/ms for in vivo neurochemical monitoring. The AAPS journal, 19: 1284–1293, 2017.

56. J. S. Farrell, R. Colangeli, A. Dong, A. G. George, K. Addo-Osafo, P. J. Kingsley, et al. In vivo endocannabinoid dynamics at the timescale of physiological and pathological neural activity. Neuron, 109. e2394:2398–2403, 2021.

57. C. Charalambous, M. Lapka, T. Havlickova, K. Syslova, and M. Sustkova-Fiserova. Alterations in rat accumbens dopamine, endocannabinoids and gaba content during win55, 212-2 treatment: The role of ghrelin. International Journal of Molecular Sciences, 22:210, 2020.

58. P. J. McLaughlin, G. A. Thakur, V. K. Vemuri, E. D. McClure,C. M. Brown, K. M. Winston, et al. Behavioral effects of the novel potent cannabinoid cb1 agonist am 4054. Pharmacology Biochemistry and Behavior, 109: 16–22, 2013.

59. A. Zimmer, A. M. Zimmer, A. G. Hohmann, M. Herkenham, and T. I. Bonner. Increased mortality, hypoactivity, and hypoalgesia in cannabinoid cb1 receptor knockout mice. Proceedings of the National Academy of Sciences, 96:5780–5785, 1999.

60. R. Tremblay, S. Lee, and B. Rudy. Gabaergic interneurons in the neocortex: from cellular properties to circuits. Neuron, 91: 260–292, 2016.

61. M. Galarreta and S. Hestrin. Frequency-dependent synaptic depression and the balance of excitation and inhibition in the neocortex. Nature neuroscience, 1: 587–594, 1998.

62. O. Friard and M. Gamba. Boris: a free, versatile opensource event-logging software for video/audio coding and live observations. Methods in ecology and evolution, 7: 1325–1330, 2016.

