## Supplemental Files for "Behavioral Decoding Reveals Cortical Endocannabinoid Potentiation during Δ^9^-THC Impairment"

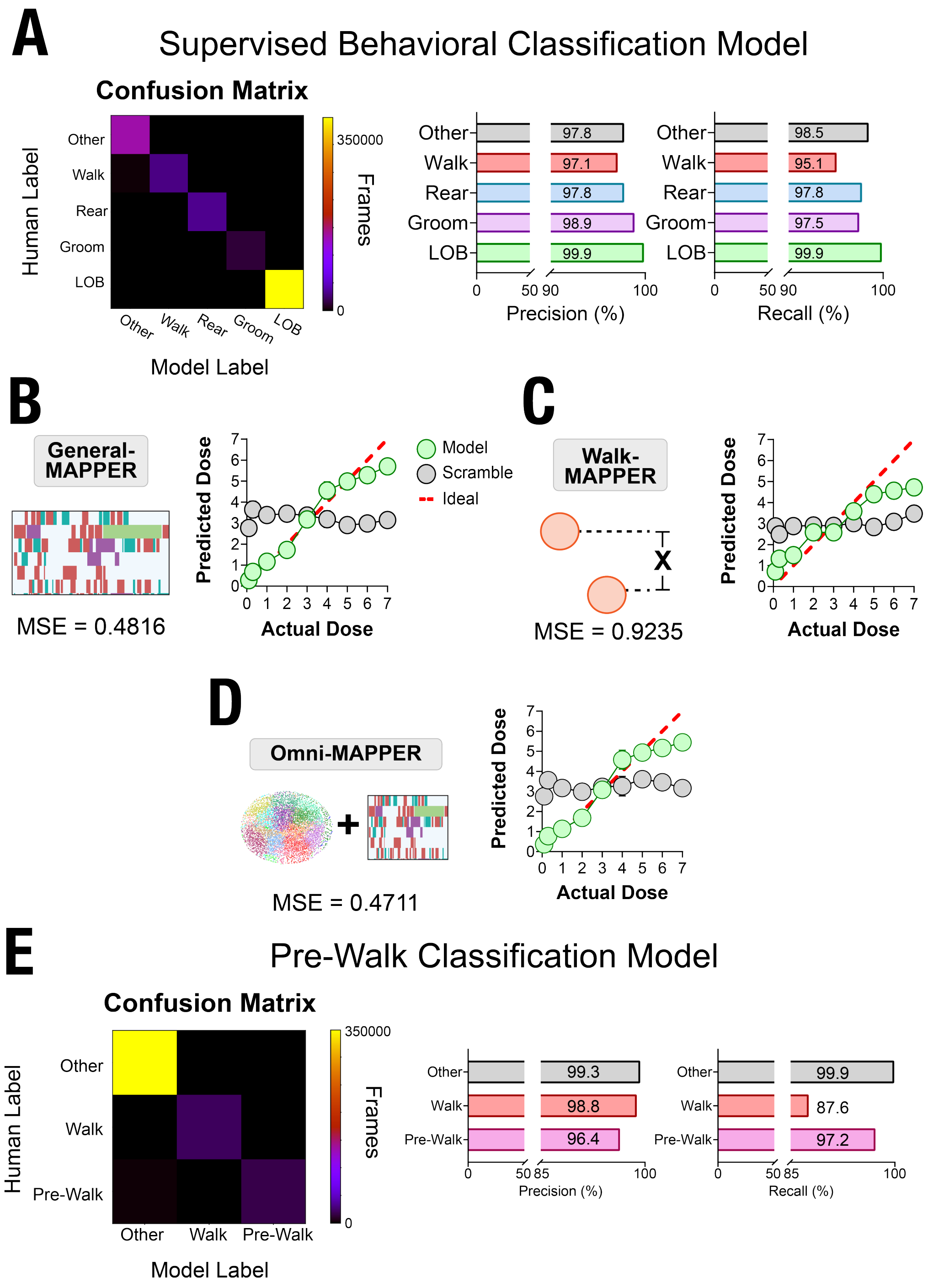


**Supplemental Figure 1. AI model and algorithm performance metrics: A)** Supervised classification of core behaviors (walking, rearing, grooming, and LOB) of animals in the linear track. Random Forest classification algorithm was trained on 176 videos of animals treated with vehicle or THC (0.1-10 mg/kg) for a total of 1.6 million frames. **B-D)** Dose prediction RF regression algorithms were assessed for their MSE scores and plotted on a test dataset to showcase model accuracy of the Core Prediction Model (**B**), Kinematic Prediction model (**C**), and the Comprehensive Prediction model (**D**). **E)** Supervised classification of pre-walk and walk behaviors of animals in the linear track for closed-loop stimulation. Random Forest classification algorithm was trained on 176 videos of animals treated with vehicle or THC (0.1-10 mg/kg) for a total of 1.6 million frames.

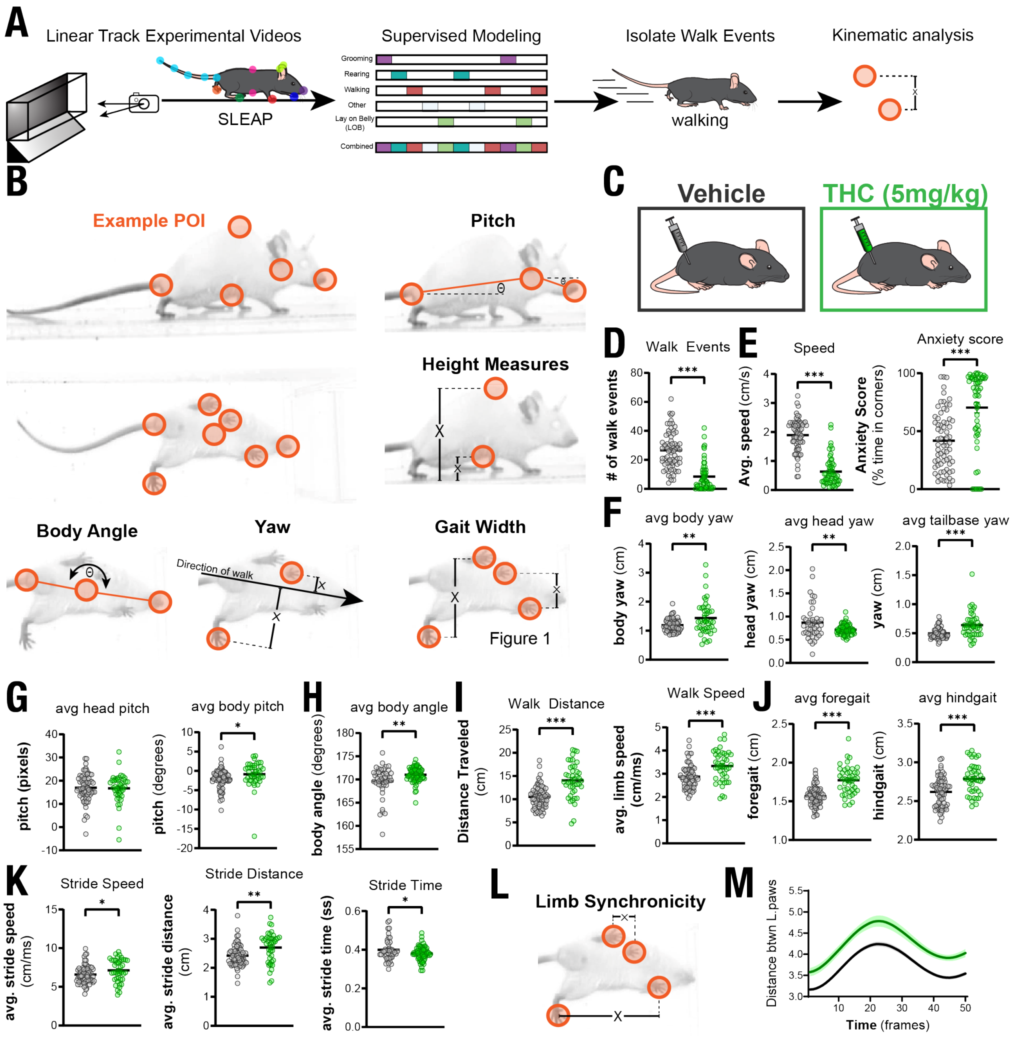


**Supplemental Figure 2. Calculating kinematic features during walking behavior: A)** Schematic showing the application of the supervised RF algorithm to identify walk events followed by latent space calculation of points of interest over the walk event. **B)** Example images of how kinematic calculations were made from points of interest across the side and bottom view from the linear track. **C)** Animals in example here were treated with either vehicle or THC at 5mg/kg and had all of the following metrics calculated across all of their walk events over a 15 min session in the linear track: **D)** Number of total walk events, **E)** Average walk speed and anxiety score, **F)** Average yaw metrics (lateral displacement along the path of movement) for the body, head, and tailbase, **G)** Average pitch of the head and nose, **H)** Average body anlge, **I)** Average distance traveled and speed during a walk, **J)** average gait width of forepaws and hindpaws, **K)** average stride speed, distance, and time. **L)** Schematic of limb synchronicity tracked over the behavioral paradigm. **M)** distance between forepay and hindpaw on the left side for the 0.5 s post walk initiation. Student's unpaired T-test, no multiplicity post-tests. *p<0.05, **p<0.01, ***p<0.001, N=70 (vehicle) or 53 (THC 5mg/kg).

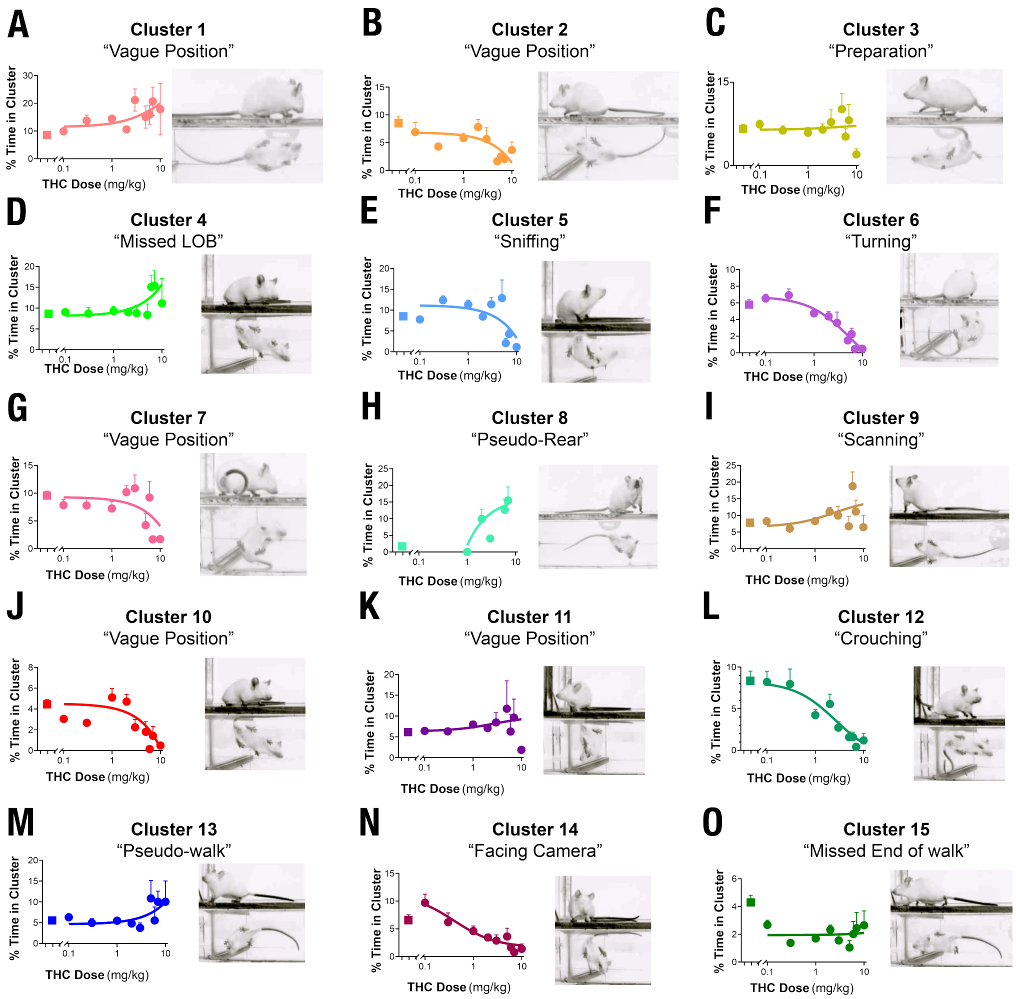


**Supplemental Figure 3. Unsupervised clustered behaviors with visual examples:** Frequency of all 15 behaviors that were isolated from 29-dimensional K-means clustering after treatment with vehicle or increasing doses of THC (0.1, 0.3, 1, 2, 3, 4, 5, 6, 7, and 10 mg/kg). Each behavior was given a name after visual inspection: **A)** Cluster 1: Vague Position, **B)** Cluster 2: Vague Position, **C)** Cluster 3: Preparation, **D)** Cluster 4: Missed LOB, **E)** Cluster 5: Sniffing, **F)** Cluster 6: Turning, **G)** Cluster 7: Vague Position, **H)** Cluster 8: Pseudo-Rear, **I)** Cluster 9: Scanning, **J)** Cluster 10: Vague Position, **K)** Cluster 11: Vague Position, **L)** Cluster 12: Crouching, **M)** Cluster 13: Pseudo-Walk, **N)** Cluster 14: Facing Camera, and **O)** Cluster 15: Missed End of walk.

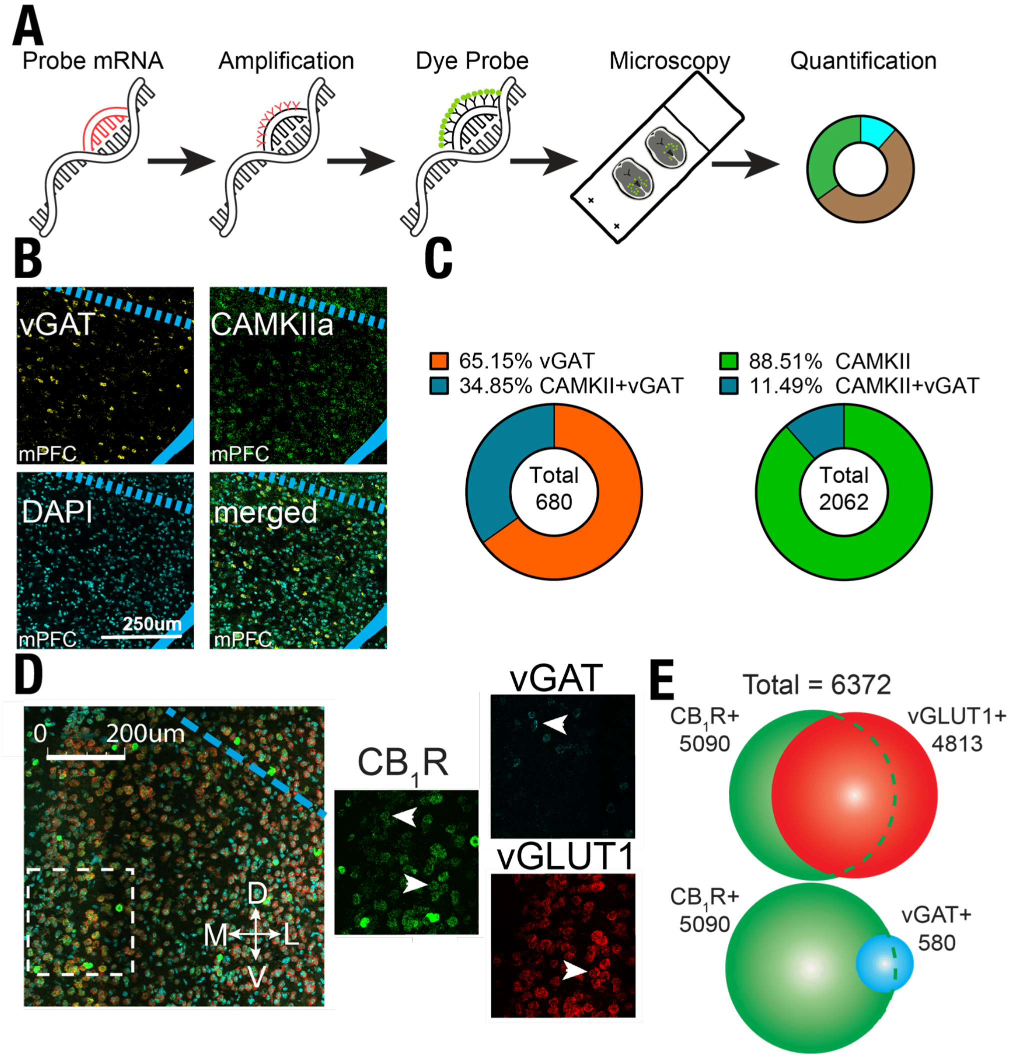


**Supplemental Figure 4. *in situ* Hybridization analysis of cell-specific $CB_1$R expression in the mPFC: A)** Schematic of RNAscope (in situ hybridization) procedure utilized throughout paper. **B)** Confocal images of WT brains targeting vGAT neurons and CAMKII neurons, stained with DAPI for neuronal selection. **C)** Quantification of co-expression of mRNA for GABAergic neurons and CAMKII neurons in the mPFC. **D)** RNAscope analysis of CB1R, vGLUT1, and vGAT mRNA across mPFC neurons. **E)** Quantification of RNAscope from D showing overlapped expression of CB1R with vGLUT1+ and vGAT+ neurons.

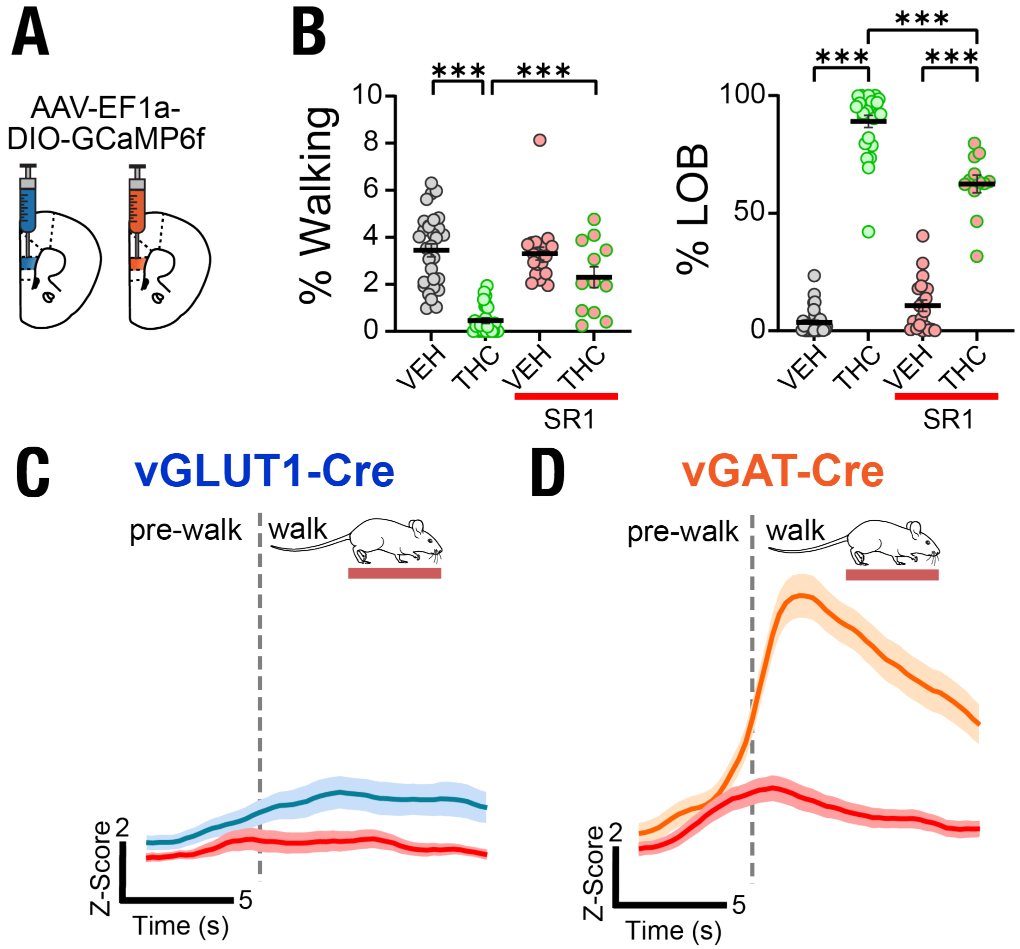


**Supplemental Figure 5. $CB_1$R Antagonism blockade of THC-induced cannabimimetic behaviors and mPFC transient activity:** **A)** Schematic of mPFC$^{vGLUT1\text{-}GCaMP6f}$ and mPFC$^{vGAT\text{-}GCaMP6f}$ for fiber photometric recordings during behavior. **B)** Percent time of walking and laying on belly after treatment with vehicle or THC (5mg/kg) alone or with a pre-treatment with SR1 (1mg/kg). **C-D)** Fiber photometry signal of glutamatergic (**C**) and GABAergic (**D**) neuron activity at walk initiation after either THC treatment (5mg/kg) or a pre-treatment of SR1 (1mg/kg) followed by a THC treatment (5mg/kg). One-Way ANOVA, Sidak post-test. ***p<0.001.

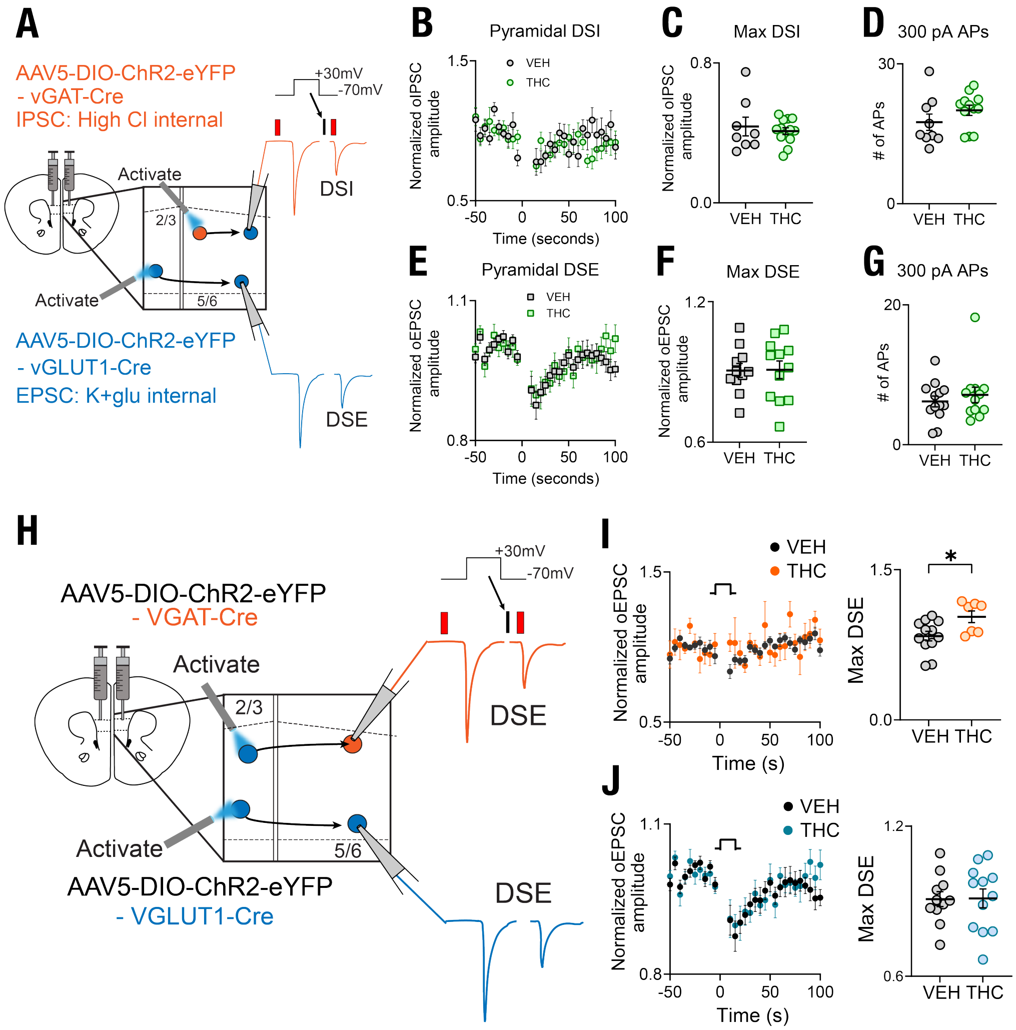


**Supplemental Figure 6. Electrophysiological investigation of THC's effects on mPFC circuitry: A)** Schematic of ex vivo electrophysiological approach to record DSE and DSI in vehicle and THC bath mPFC tissue samples. **B)** Normalized oIPSC after vehicle or THC bath application. **C)** Maximum DSI from normalized oIPSC amplitude. **D)** Number of action potentials after 300 pA stimulation. **E)** Normalized oEPSC after vehicle or THC bath application. **F)** Maximum DSI from normalized oEPSC amplitude. **G)** Number of action potentials after 300 pA stimulation. **H)** Schematic for paired pulse ratio from glutamatergic and GABAergic neurons. **I)** vGLUT1 Paired Pulse Ratio. **J)** vGAT Paired Pulse Ratio. Student's T-test, no multiplicity post-tests. *p<0.05.

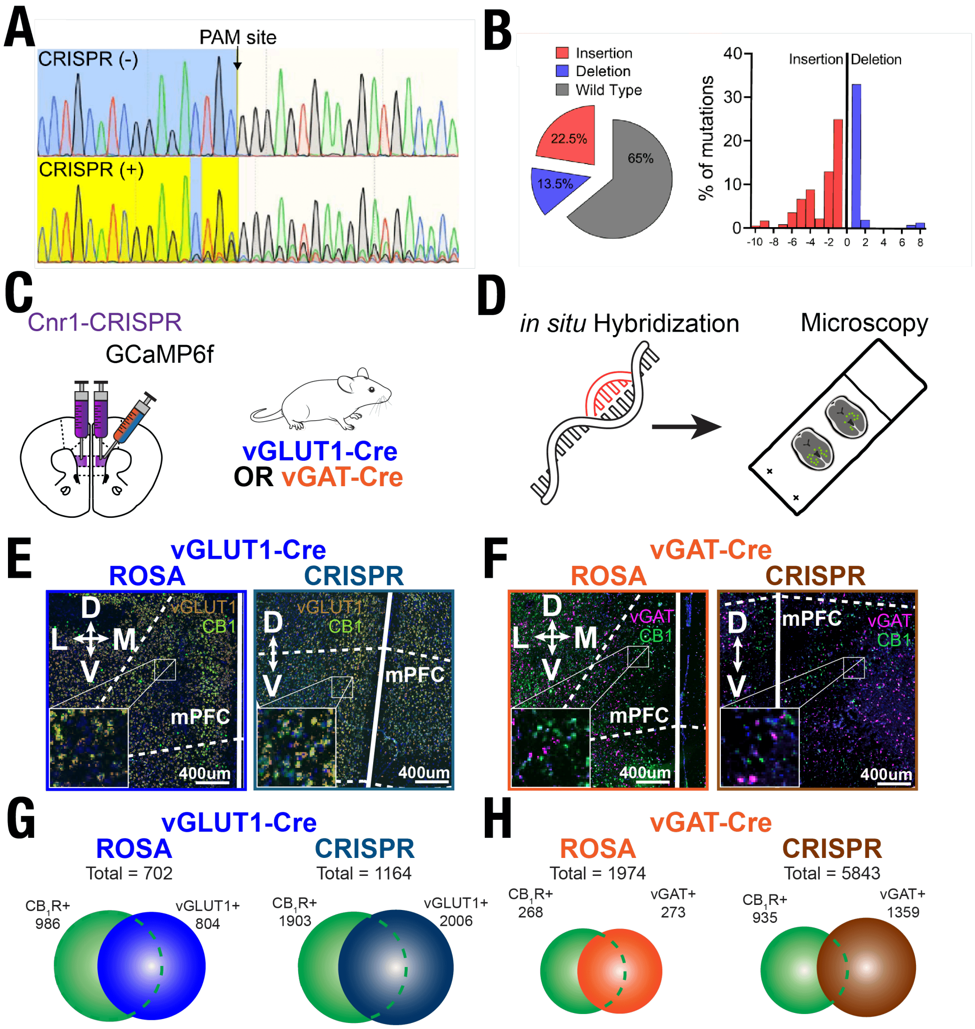


**Supplemental Figure 7. CB$_1$R CRISPR validation of knockout in mPFC neurons: A-B)** Fluorescent-Activate Cell Sorting (FACS) of mPFC vGAT neurons treated with AAV-SaCas9-sgCnr1 showing the point of mutation (**A**) and the quantification of insertions and deletions (**B**). **C)** Schematic of bi-lateral injection of AAV-SaCas9-sgSnr1 into mPFC of vGLUT-Cre and vGAT-Cre animals. **D)** Simple schematic of *in situ* hybdirization procedure from Supplementary Fugyre S4. **E-F)** Confocal images of mPFC$^{vGLUT\text{-}CRISPR}$, mPFC$^{vGLUT1\text{-}ROSA}$, mPFC$^{vGAT\text{-}CRISPR}$, and mPFC$^{vGAT\text{-}ROSA}$ neurons. **G-H)** Quantification of Cnr1 Knockout in mPFC$^{vGLUT\text{-}CRISPR}$ and mPFC$^{vGAT\text{-}CRISPR}$ neurons across experimental animals.
